# Convergent IGHV3-53/3-66 antibodies elicited by Omicron BA.1 infection broadly neutralize emerging SARS-CoV-2 variants

**DOI:** 10.64898/2026.08.15.745004

**Authors:** Naveenchandra Suryadevara, Seth J. Zost, John M. Powers, Bernadeta Dadonaite, Pavlo Gilchuk, Elad Binshtein, Suzanne Scheaffer, Sarah R Leist, Luke Myers, Silvia Ravera, Lily E. Adams, Laura S. Handal, Shruthi Kannan, Edgar Davidson, Benjamin J. Doranz, Andrew Trivette, Masako Abney, Doan C. Nguyen, F. Eun-Hyung Lee, Robert H. Carnahan, Jesse D. Bloom, Ralph S. Baric, Michael S. Diamond, James E. Crowe

## Abstract

Natural SARS-CoV-2 infections or vaccinations induce neutralizing antibodies (nAbs) offer protection from severe disease. The shared use of *IGHV3-53/3-66* genes makes this class of monoclonal antibodies (mAbs) a public clonotype and is well established, but the evolution and structural basis of how these public antibodies maintain broad binding and acquire potent neutralizing activity is not completely understood. To understand how these features are facilitated by the *IGHV3-53/3-66* germline segments and enhanced by somatic mutations, we investigated the biology of a panel of 242 human mAbs isolated from an individual infected with SARS-CoV-2 BA.1 strain and recovered. Interestingly, a mAb designated COV2-3731 encoded by *IGHV3-53/IGKV1-33* retained potent neutralizing activity against SARS-CoV-2 variants BA.2.86, JN.1, KP.2, BA.3.2, and, to some extent, KP.3. Studies using deep mutational scanning with a BA.2 lentiviral library and determination of the structural complex of the BA.2 S protein and COV2-3731 Fab fragments using cryo-EM revealed key contact residues. Further, germline revertant analysis of the COV2-3731 mAb provided additional insights into how this COV2-3731 and other *IGHV3-53/3-66*-encoded public antibodies evolve to gain breadth against antigenically distinct SARS-CoV-2 variants such as BA.2.86, JN.1, KP.2, and BA.3.2.

## Introduction

The coronavirus 2019 (COVID-19) pandemic, caused by severe acute respiratory syndrome coronavirus 2 (SARS-CoV-2), has substantially reshaped the human antibody repertoire, and understanding antibody evolution is important to design new vaccines. The trimeric spike (S) glycoprotein, facilitates attachment to angiotensin-converting enzyme 2 (ACE2), the host receptor, which is the primary target of neutralizing antibodies (nAbs) and forms the basis for nearly all licensed therapeutics^1–4^. Antibodies generated against the receptor-binding domain (RBD) dominate in offering protection, as they block ACE2 from engaging with spike protein, thereby averting viral entry^5–8^.

Although human B-cell repertoire is so diversified, detailed studies using several convalescent donors responses to SARS-CoV-2 infection exposed shared antibody gene usages and similar structural motifs, revealing convergent evolution of B-cell lineages ^9–13^. Amongst these lineages, *IGHV3-53* and *IGHV3-66* germline genes encoded antibodies have emerged as one of the most prevalent and genetically identified to be “public” clonotypes in COVID-19 infection. Moreover, these two heavy-chain variable genes diverge by only a single amino-acid residue in their framework 1 region and are functionally interchangeable, yielding similar or identical paratopes that engage the RBD^10,11,14^. Nonetheless, motifs such as asparagine-tyrosine (Asn-Tyr) of heavy-chain complementarity-determining region 1 (CDRH1) and serine-glycine (Ser-Gly-Gly-Ser) of CDRH2 loops are germline-encoded and are critical for recognizing the ridge of receptor-binding domain, allowing virus neutralization with nominal diversification through somatic hypermutation^15–17^.

B cell repertoire analyses to date against SARS-CoV-2 have established that *IGHV3-53/3-66*-encoded antibodies have dominated RBD-directed immunity across different individuals and as well as cohorts^10,12,18^. Large-scale datasets have well-defined “public” antibody clusters that were characterized by conventional V(D)J usage, either short or long CDRH3 loops (typically 9 to 18 residues), and with persistent light-chain pairings—most notably encoded by either *IGKV1-9* or *IGKV3-20*^19^. The combination of high-resolution structures of these antibodies, along with sequence analyses, has offered a greater understanding of human antibody evolution. For instance, Tan *et al*., delineated public clonotypes encoded by *IGHV3-53/3-66* genes: one pairing with *IGKV1-9* and the other with *IGKV3-20* encoded light chains ^19,20^. Of note, each clonotype was identified to carry a distinctive sequence in CDRH3 that facilitates complementarity of specific heavy- and light-chains. Furthermore, it was noted that the choice of light-chain dictated subtle changes in RBD engagement through Π-Π (pi-pi) stacking and electrostatic interactions, on the other hand, somatic mutation such as Y58F in CDRH2 enhanced affinity by improving the hydrogen-bond network at the interfaces of epitope-paratope ^20^. Together, these results established that antibodies encoded by *IGHV3-53/3-66* as a germline “public framework” that can be refined by restricted maturation to attain broad and potent neutralization^11,19,21^.

*IGHV3-53/3-66*-encoded antibody evolution exemplifies the requirement for the recruitment of existing germline genes and their adaptation by somatic hypermutations. Early in infection, antibodies encoded by *IGHV3-53/3-66* appeared with few substitutions, offering immediate neutralization capability^9,22^. Over time, it was noted that through affinity maturation and targeted substitutions, such as Y58F or S31R, boosted their capacity to bind broadly and potency ^19,23^. Nonetheless, the buildup of mutations in circulating viral variants has repetitively tested the *IGHV3-53/3-66*-encoded class of antibodies. Likewise, 15 substitutions in RBD region have led to the emergence of the Omicron variants, resulting in prevalent escape from germline *IGHV3-53/3-66*-encoded nAbs^15,20,24–27^.

Further, Fan *et al.,* described ConBA-998, a mAb encoded by *IGHV3-66* that was isolated from an unvaccinated individual following primary Omicron BA.1 infection^28^. Although ConBA-998 potently neutralized BA.1 (half-maximal inhibitory concentration [IC ] value 0.003 µg mL ¹), it failed to bind the ancestral S protein, defining it as an Omicron-specific nAb. A cryo-electron microscopy structure at 3.4 Å resolution revealed a binding angle distinct from that of canonical *IGHV3-53/3-66*-encoded nAbs and also demonstrated that ConBA-998 triggers S1-subunit shedding via remodeling at the paratope interface-a mechanism reported to be exhibited by CB6 and P2C-1F11 antibodies ^28^. Above observations emphasize that even a conserved germline framework might allow structural flexibility, enabling adaptation to different viral mutations in those interacting epitopes, extending the evolutionary potential of this public clonotype^28^. Furthermore, recent studies demonstrate lack of ancestral imprinting promotes infections with recent variants such as BA.3.2.2 in children^29,30^.

Taken together, these studies underscore the importance of public clonotypes encoded by IGHV*3-53/3-66* germline genes, which provide a classic scaffold enabling recognition of a viral protein and led to the evolution of lineages to accommodate the emergence of viral variants. The favorable structural assembly of these antibodies confers potent neutralization against SARS-CoV-2. Yet, this lineage diversified as the virus evolved, achieving new paratope conformations and alternating functional mechanisms. Repertoire studies of individuals with SARS-CoV-2 infections or vaccinations identified consistent usage of *IGHV3-53* gene, suggesting that germline genes remain a privileged entry point for recognition of RBD across different variants. Understanding the molecular and evolutionary trajectory of the antibodies encoded by *IGHV3-53* is critical for designing new-generation vaccines. Insights from convergent/public antibodies evolution not only reveal how the immune system repeatedly solves the problem of neutralizing the virus but also offer a blueprint for the principles underlying engineering broad and variant-resistant immunity ^5,21,28,31^.

## Results

### Enrichment of ACE2-blocking B cells and screening for antibody reactivity

To isolate SARS-CoV-2 Omicron (BA.1 antigen) specific B cells, PBMCs collected from a BA.1-convalescent donor three months after infection were enriched for Pan B cells and stained with phenotypic markers, analyzed by flow cytometry. Of enriched pan-B cells, approximately 40% of total cells fell within the lymphocyte gate, and 92% of these exhibited the expected forward– side scatter profile of viable single cells (**Figure 1A**). These single cells were further gated for IgD/IgM and CD19 cells. Within CD19^+^compartment, 1.02% of cells were double-positive for ACE2 and BA.1 RBD, representing antigen-specific B cells, while 0.3% of cells exhibited an ACE2-blocking phenotype (**Figure 1A**). Populations of B cells, either ACE2-blocked or ACE2-positive, were sorted, plated on 3T3 cell culture monolayers that display CD154 and secrete IL-21 and B-cell activating factor (BAFF), and allowed them to expand for 7 to 9 days to differentiate into antibody-secreting cells (ASCs). On day 7, the supernatants were screened by ELISA for antibody reactivity and function. ASC-derived antibodies exhibited avid binding to BA.1 RBD, (**Figure 1B**). Antibodies from the supernatants of cells sorted by blocking ACE2 blocked the ACE2–RBD interaction, reducing ACE2 binding by more than 90% in a dose-dependent manner, confirming that the ACE2-blocking enrichment strategy is likely effective. In contrast, supernatants from non-ACE2-blocking ASCs showed negligible inhibition (**Figure 1C**). Overall, our results confirmed that the successful isolation of ACE2-blocking memory B cells that produce antibodies might be capable of restricting the viral receptor interaction.

**Figure 1.**
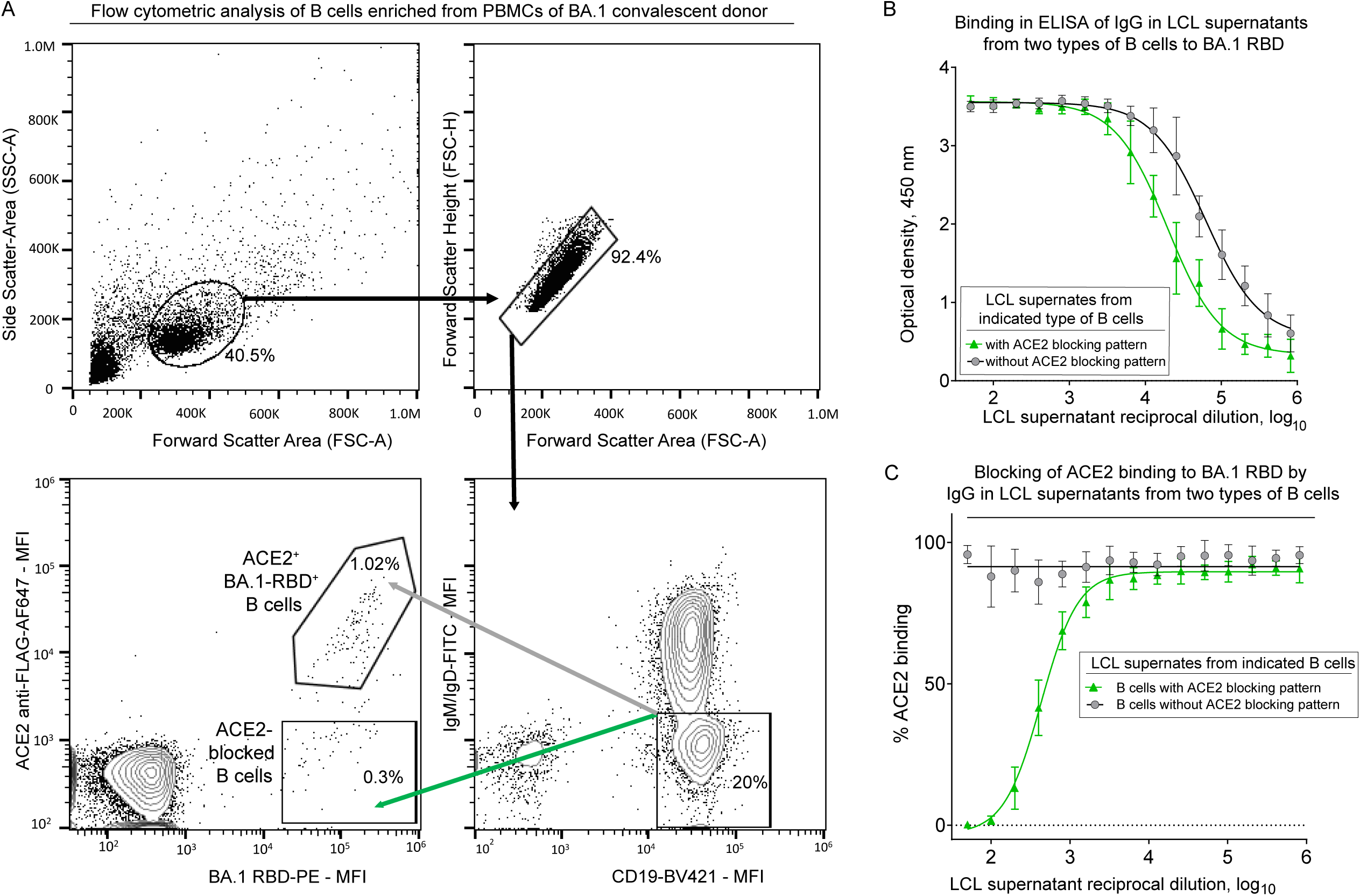
Enrichment of ACE2 blocking B cells and supernatant antibody reactivity. **A.** Flow cytometry of memory B cells in total B cells enriched by negative selection using magnetic beads; cells were stained with anti-CD19 antibody conjugated to Brilliant Violet 421(BV421) and anti-IgM and anti-IgD antibodies conjugated to fluorescein isothiocyanate (FITC), BA.1 receptor binding domain (RBD), and human angiotensin-converting enzyme 2 with FLAG tag (hACE2-FLAG). Plots show single cells, CD19+IgD–IgM− population gating, cells labeled with biotinylated RBD were detected using phycoerythrin (PE)-conjugated streptavidin, and cells labeled with hACE2 flag tag were detected with anti-FLAG-Alexa fluor 647 (AF647) antigens. **B -C.** Antibody-secreting cell (ASC) supernatant antibody reactivity. Cells after flow sorting BA.1 RBD^+^ ACE2^+^ (double positive) and BA.1 RBD^+^ and ACE2^-^ (B cells that blocked ACE2 from binding) were stimulated in 10 wells *in vitro* on feeder layers expressing CD40L and secreting IL-21 and BAFF. The supernatants were tested in a fifteen-point dilution series to check for BA.1 RBD binding and ACE2 blocking from binding to BA.1 RBD. **B.** Binding of antibodies in supernatants from BA.1 RBD^+^ ACE2^+^ (shown in a grey circle with a black line, while BA.1 RBD^+^ and ACE2^-^ (shown in triangles with green line). **C.** Binding of ACE2 to BA.1 RBD BA.1 RBD^+^ ACE2^+^ (shown in a grey circle with a black line, while BA.1 RBD^+^ and ACE2^-^ (shown in triangles with green line).

### Virus neutralization, and competition-binding analyses

A total of nearly 300 paired heavy- and light-chain variable gene sequences were recovered from two different RNA-seq workflows (using Bruker Beacon or 10x Genomics technologies), as described in detail in the methods section. The detailed gene usage and accession numbers of those mAb sequences are provided in **Table S1**. To evaluate the antigenic breadth of the antibodies, we expressed and purified mAbs at the microscale level as described previously^32,33^ and screened for binding against SARS-CoV-2 and SARS-CoV antigens. Most antibodies bound strongly to SARS-CoV-2 S6P_ecto_, RBDs, including WT (Wuhan-Hu-1) and BA.1, with limited cross-reactivity to SARS-CoV S2P_ecto,_ and RBD (**Figure 2A, Figure S1**). Binding to the S ectodomain protein correlated with RBD binding, demonstrating that the antibodies target conformationally accessible epitopes on the trimeric S protein. Use of pseudotyped viruses for neutralization revealed that a substantial number of antibodies neutralized the ancestral Wuhan-Hu-1 D614G strain, and around 25 -30% of them had neutralization activity against BA.1.1 lineage (**Figure 2B**). While a subset of the antibodies preserved robust neutralization at the highest tested dilutions, suggesting they target conserved RBD regions **(Figure S2 & S3)**. Further, competition-binding assay using the reference mAbs rLY-CoV1404 (Class 3 site), rS2K146 (Class 1 & 2 site), and rCR3022 (Class 4 site) revealed multiple distinct epitope binding patterns (**Figure 2C**). A significant fraction of antibodies competed with rLY-CoV1404, indicating recognition of the receptor-binding motif (RBM), whereas others overlapped with Class 4 site as rCR3022, mapping to a cryptic and more conserved, region. Together, these data explain that the BA.1 convalescent B cell repertoire encodes a diverse collection of RBD-targeting antibodies, some of which retain cross-neutralizing activity against different Omicron sublineages.

**Figure 2.**
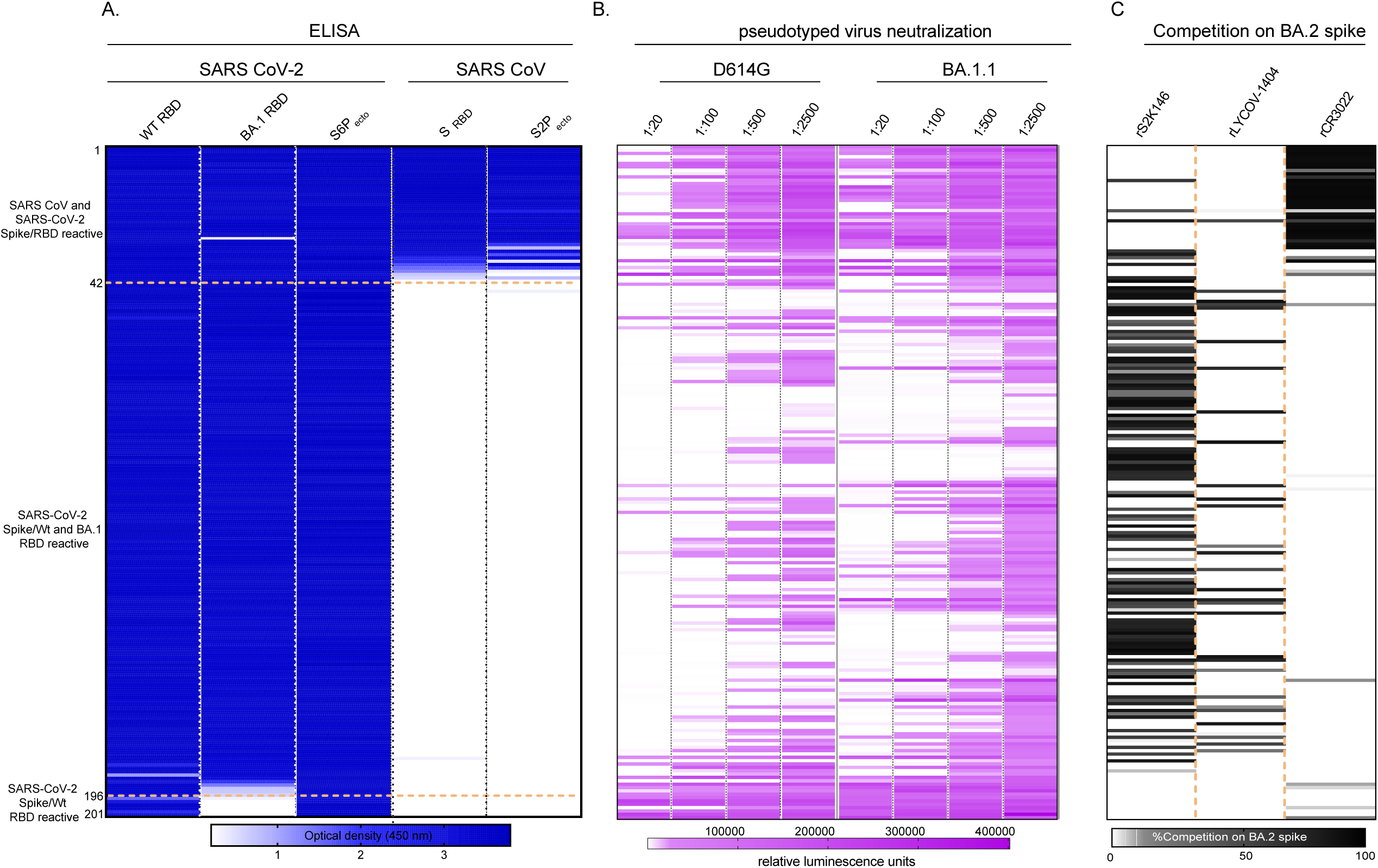
Microscale-expressed mAb binding, pseudotyped virus neutralization, and competition-binding analysis. **A.** MAb specificity or reactivity for each of the 2 S proteins or subdomains. Heatmap shows binding of 201 mAbs expressed recombinantly, representing optical (OD) values collected at 450 nm for each antigen (range, 0.5 to 4.0). White indicates a lack of detectable binding, blue indicates binding, and darker blue indicates higher OD values. **B.** Microscale-expressed mAb neutralization using pseudotyped viruses. White indicates neutralization, pink indicates moderate, and darker red indicates no neutralization. **C.** Competition of microscale-expressed mAbs using BA.2 S protein and reference mAbs rS2K146, rLyCOV-1404, rCR3022. The binding of reference mAbs to BA.2 S protein was measured in the presence of saturating competitor mAbs in a competition ELISA and normalized to binding in the presence of the isotype-matched negative control mAb DENV r2D22. Black, full competition (<25% binding of reference antibody); grey, partial competition (25–60% binding of reference antibody); white, no competition (>60% binding of reference antibody).

### Neutralization breadth, epitope mapping, and mutational sensitivity of mAbs

We evaluated the breadth of RBD-directed mAbs against a wide range of pseudotyped and authentic SARS-CoV-2 variant viruses. Mabs, namely, COV2-3600, COV2-3605, COV2-3685, COV2-3703, and COV2-3731, confirmed potent neutralization against multiple variants, with a low nanogram per milliliter (ng/mL) range of half-maximal inhibitory concentration (IC□□) values against early isolates and Omicron sublineages (**Figure S2 & S3)**. Whereas COV2-3619 and COV2-3678 retained moderate potency against heavily mutated Omicron subvariants, including BA.2.75.2, XBB.1.5, and KP.2 (**Figure 3A, Figure S2 & S3)**. Notably, certain mAbs like COV2-3600 and COV2-3731 maintained activity even against the highly immune-evasive variants JN.1, KP.3 and most recent variant BA.3.2 suggesting engagement of structurally conserved epitopes.

**Figure 3.**
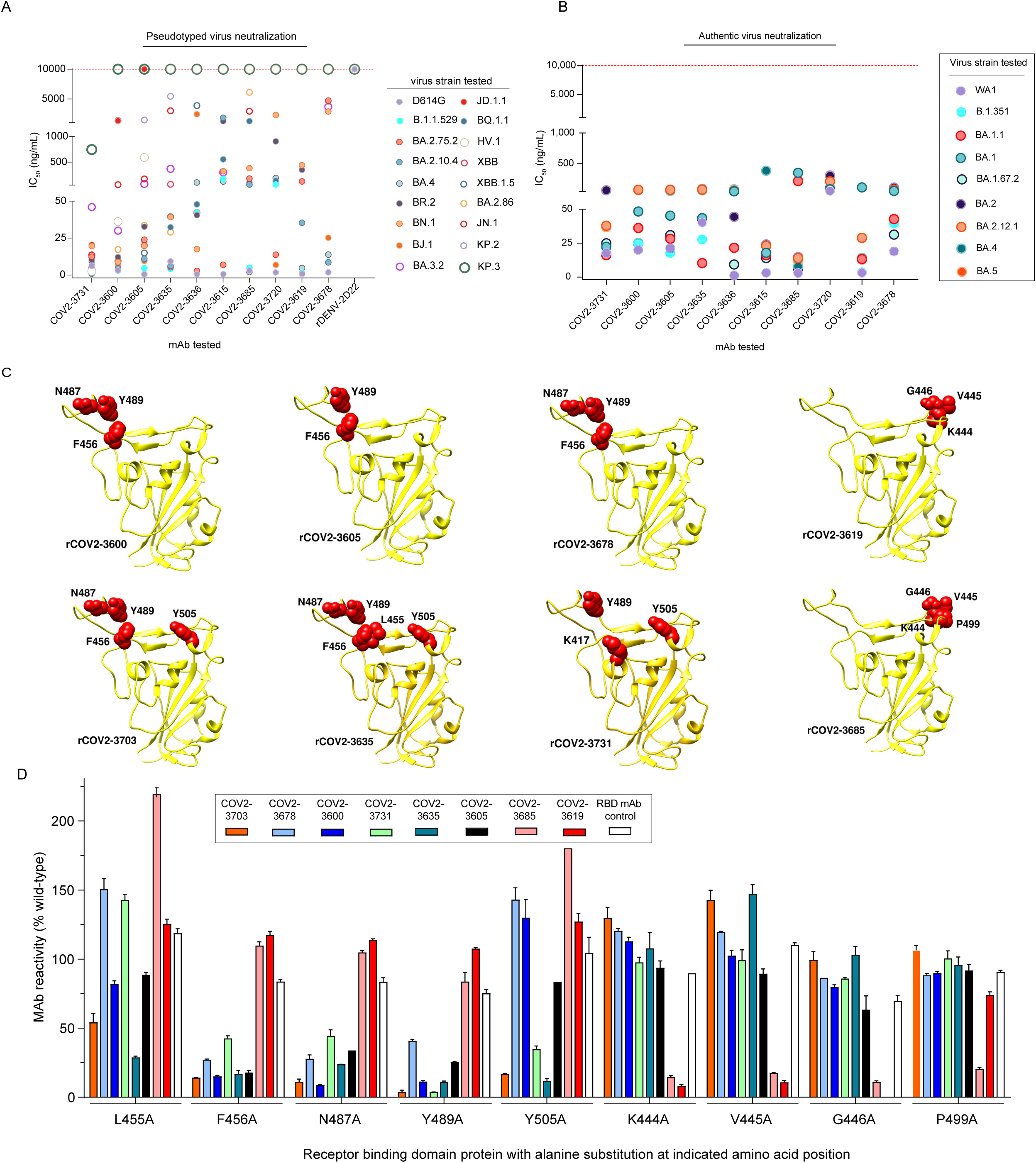
Neutralization breadth, epitope mapping, and mutational sensitivity of SARS-CoV-2 RBD-specific mAbs. **A.** Pseudotyped virus neutralization by representative mAbs against diverse SARS-CoV-2 variants. IC_50_ values were determined against pseudotyped viruses bearing S protein mutations corresponding to D614G, B.1.1.529, BA.1.1, BA.2.75.2, XBB, XBB.1.5, BA.4, BR.2, BN.1, JN.1, BJ.1, KP.3, and BA.3.2 variants. Neutralization assays were performed in technical duplicates with at least two independent experimental replicates. **B.** Authentic virus neutralization by the same panel of mAbs tested against SARS-CoV-2 variants WA1, B.1.351, BA.1, BA.1.1, BA.1.617.2, BA.2, BA.2.12.1, BA.4, and BA.5. Each data point represents an individual IC_50_ measurement; dashed red line denotes assay upper detection limit. Neutralization assays were performed in technical duplicates with at least two independent experimental replicates. **C.** Structural mapping of key RBD residues recognized by representative neutralizing mAbs. Antibody-specific contact residues identified by alanine scanning mutagenesis are shown in red on the SARS-CoV-2 RBD structure (PDB: 6XCN). **D.** Mutational sensitivity profiling of mAb binding to RBD alanine mutants. Bars indicate the relative binding of each mAb (normalized to WT RBD, set to 100%). Colors correspond to individual antibodies as indicated in the legend; the blue bar denotes the RBD mAb control. Error bars represent mean ± range of technical replicates.

Using alanine scanning mutagenesis as described previously,^34^ we were able to map epitopes of certain representative antibodies, which correlated with their binding, neutralization, and competition. Antibodies like COV2-3600, COV2-3605, and COV2-3731 targeted class I site of receptor-binding domain, on the ridge and overlapped with the ACE2-binding interface by involving critical residues like F456, Y489, N487, and Y505 (**Figure 3B**). On the other hand, mAbs COV2-3619 and COV2-3685 recognized residues such as K444, G446, and P499, an epitope typical of Class 3 antibody binding. Notably, class-I competing mAb COV2-3731 presented a diverse binding footprint that involved residue K417, which was consistent with mAbs that cross-neutralize Omicron and Omicron variants (**Figure 3C**). Together, our findings were consistent with functional properties of mAbs reported previously by other groups and identified more somatically mutated antibodies that exhibit activity against variants of concern (VOCs).

### Structural mapping of the binding interface of COV2-3731 and related antibodies on SARS-CoV-2 RBD

To understand the structural basis of COV2-3731 neutralization breadth, we next executed epitope mapping by deep mutational scanning (DMS) studies along with cryo-electron microscopy (Cryo-EM) studies of COV2-3600, COV2-3619, and COV2-3731 Fabs that yielded high-resolution structures in complex with BA.2 spike protein (**Figure S4, S5, and S6**). Deep mutational scanning delineated that COV2-3731 neutralization is sensitive to substitutions at residues Y420, N487, and Y505, whereas COV2-3600 neutralization is compromised by mutations near residues Y420 and F496 (**Figure 4A–B**). Mapping of COV2-3619 indicated sensitivity to alterations at positions 444-445, defining a distinct but partially overlapping neutralization footprint relative to the other antibodies reported for class 3 site on RBD (**Figure 4C)**.

**Figure 4.**
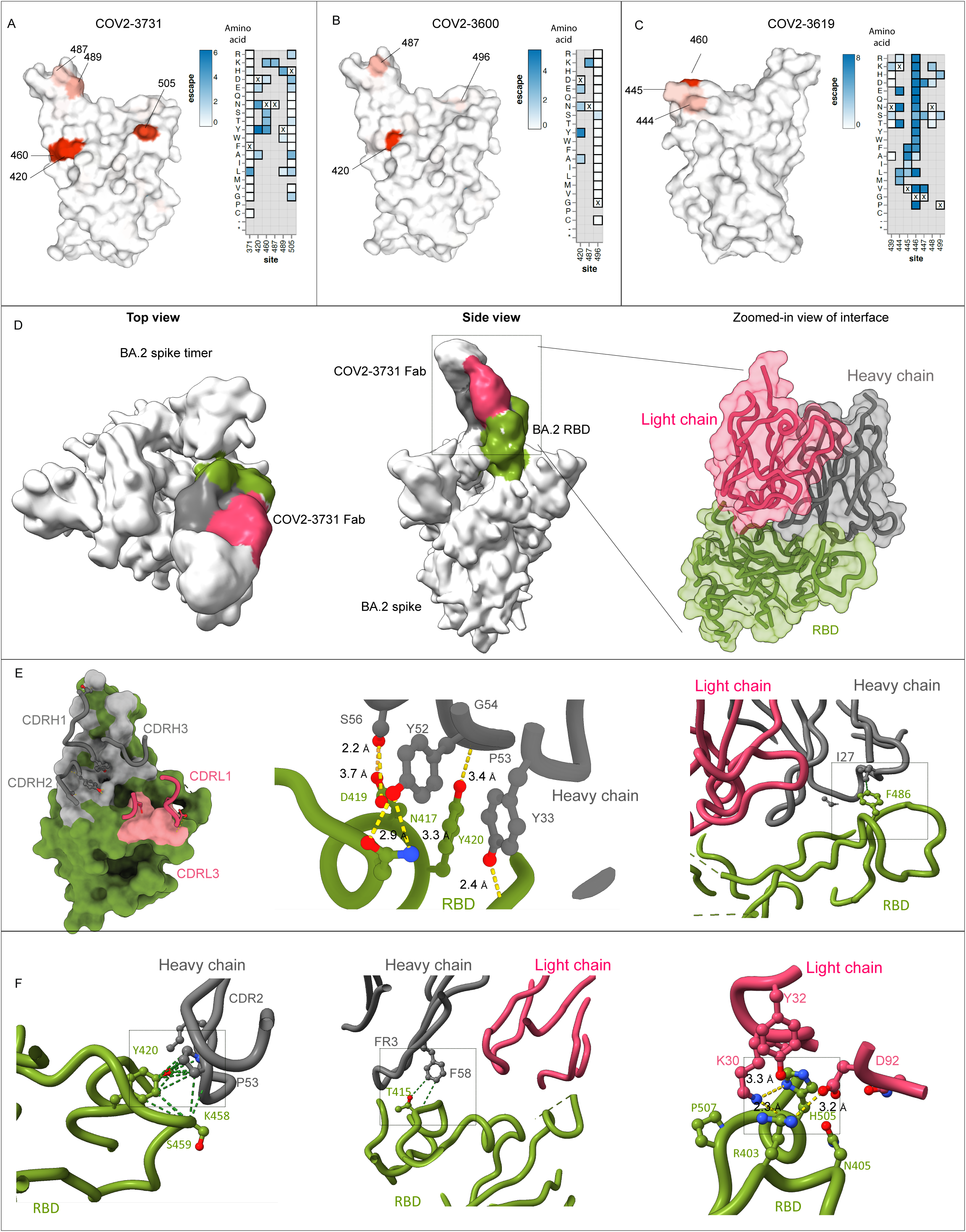
Structural mapping and binding interface of COV2-3731 and related antibodies on the SARS-CoV-2 RBD. **(A–C)** Surface representation of the RBD colored by summed neutralization escape scores at each site for antibodies COV2-3731 **(A),** COV2-3600 **(B),** and COV2-3619 **(C).** as measured by BA.2 spike deep mutational scanning. In surface representation, white indicates no escape, and darker red indicates stronger neutralization escape caused by mutations at that site for that antibody. The heat map shows escape for individual mutations, with parental amino-acid identities indicated by **x,** and gray indicating non-measured mutations. Interactive heatmaps showing neutralization escape for the full spike are at https://dms-vep.org/SARS-CoV-2_Omicron_BA.2_spike_DMS_Crowe_mAbs/COV3731_escape_plot.html for COV2-3731, https://dms-vep.org/SARS-CoV-2_Omicron_BA.2_spike_DMS_Crowe_mAbs/COV3600_escape_plot.html for COV2-3600, and https://dms-vep.org/SARS-CoV-2_Omicron_BA.2_spike_DMS_Crowe_mAbs/COV3619_escape_plot.html for COV2-3619. **(D)** Cryo-EM structure of the COV2-3731 Fab in complex with BA.2 spike trimer. The top and side views depict the Fab (gray) bound to one RBD (green) within the trimer (remaining protomers in blue and orange). The inset shows a zoomed-in view of the binding interface, with the heavy or light chains colored gray or pink, respectively, and the RBD in green. **(E)** Detailed view of the COV2-3731 paratope–epitope interactions. Left, surface representation of RBD (green) and the COV2-3731 Fab highlighting CDR loops (heavy chain, gray; light chain CDRL1 and CDRL3, pink). Middle, close-up of hydrogen-bond and van der Waals contacts between RBD residues D419, N417, and Y420 and the COV2-3731 heavy chain (distances in Å). Right, interactions between RBD residue F486 and light chain residue I27 at the interface. **(F)** Additional interface details showing interactions between COV2-3731 and the RBD. Left, contacts involving RBD residues K458 and S459 with heavy chain residues P53 and Y420. Middle, hydrogen bonds between RBD residue T415 and heavy chain residues F58 and FR3 loop. Right, polar interactions involving light chain residues Y32, K30, and D92 with RBD residues R403, N405, and H505.

Cryo-EM reconstruction of COV2-3731 Fab bound to BA.2 S protein trimer uncovered that COV2-3731 engages through complementarity-determining regions (CDRs) of both heavy (V_H_) and light (V_L_) chains at the apex of the receptor-binding ridge, making extensive contacts (**Figure 4D**), CDRH1, CDRH2, and CDRH3 of heavy chain majorly contributed to form hydrogen bonds and van der Waals interactions with RBD residues such as N417, Y420, and D419, establishing a tight interface (**Figure 4E, and S7**). Of note, light-chain CDRL1 and CDRL3 also contributed to additional contacts with residues close to the RBM.

A closer look at the interface highlighted numerous key stabilizing interactions, mainly heavy-chain residues such as Y52, P53, and S56 forming a hydrophilic pocket near residue Y420, whereas light-chain residues like Y32 and K30 alleviate the periphery of the binding ridge (**Figure 4E–F**). Further, framework residues of COV2-3731 were noted to make stronger interactions with conserved residues like K458 and S459 of RBD, likely contributing to mutational tolerance of the newest omicron lineages.

Together, structural and functional analyses imply that COV2-3600 and COV2-3731 neutralize SARS-CoV-2 variants predominantly through high-affinity engagement with RBD and occupying the conserved residues that are necessary for ACE2 binding (**Figure 4D, and S7)**. COV2-3731 primary interactions are focused on N417 and Y420 residues, which are receptor-binding ridge structural core. In parallel, the COV2-3731 Fab light chain formed superficial contacts that stabilized the epitope, describing the exceptional potency of COV2-3731 despite its limited number of somatic mutations. Overall, COV2-3600 and COV2-3731 had distinct binding footprints but they overlap for epitope suggesting convergent recognition of a conformational RBD epitope, which likely is the cause for neutralization breadth. Nonetheless, such interactions were noted to impose selective pressure, promoting substitutions outside the ACE2-binding interface, which likely accounts for the low frequency of escape mutations observed at those antibody-binding sites among circulating Omicron sublineages.

### Protective efficacy of mAbs in SARS-CoV-2 BA.5–challenged mice

Next, the protective efficacy of potent and broad mAbs such as COV2-3600 and COV2-3731 was tested *in vivo*, using BALB/c mice. Mice were prophylactically administered with selected mAbs along with rLYCoV-1404 (positive control), or rDENV-2D22 (isotype-matched negative control mAb recognizing dengue virus envelope protein) prior to intranasal challenge with SARS-CoV-2 virus. Mice receiving PBS or the control IgG rDENV-2D22 showed signs of weight loss beginning at 4 days post-infection (dpi), reaching ∼25 to 30% by 7 to 8 dpi (**Figure 5A**). Whereas mice that received mAbs COV2-3600, COV2-3731, and rLYCoV-1404 (positive control) did not exhibit any weight loss, further conferring complete protection from SARS-CoV-2 challenge. Consistently, survival analysis confirmed rapid mortality in the rDENV-2D22 (isotype-matched antibody control) cohorts by 7 to 8 dpi, whereas mice from all other antibody-administration survived (**Figure 5B**).

**Figure 5.**
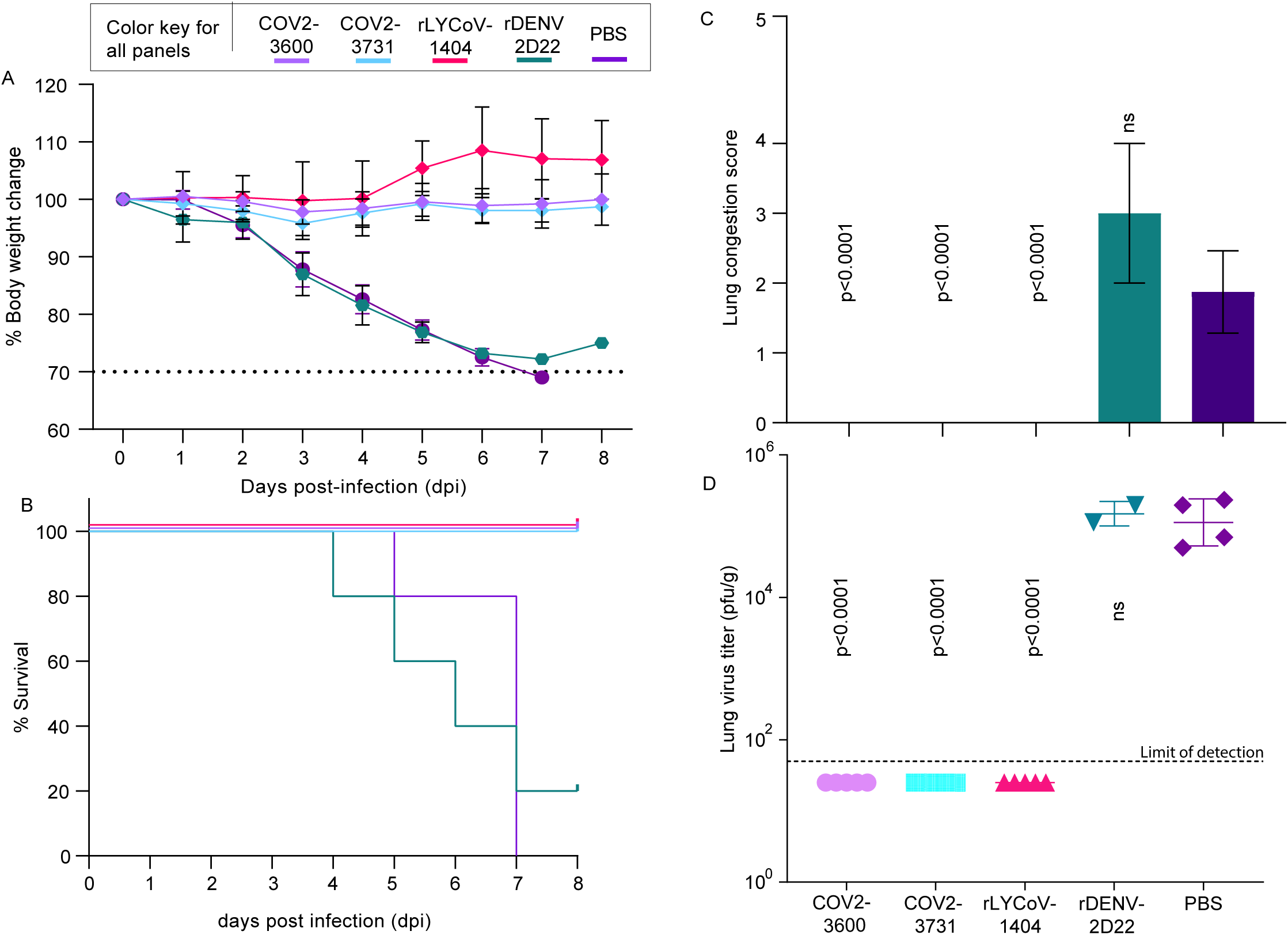
Protective efficacy of mAbs in SARS-CoV-2 BA.5–challenged mice. **(A)** Body weight change of female BALB/c mice following intranasal challenge with 10^5^ PFU SARS-CoV-2 BA.5 MA. Mice were treated with 10 mg/kg of rCOV2-3600, rCOV2-3731, or control antibodies (rLY-CoV1404, rDENV-2D22) one day before infection. Weight was monitored daily and plotted as a percentage change relative to baseline. The dotted line indicates the humane endpoint threshold (70% of the starting weight). Data represent mean ± SEM. **(B)** Kaplan–Meier survival curves for the same groups shown in (A). Mice were monitored for 8 days post-infection (dpi); survival differences were analyzed by the log-rank (Mantel–Cox) test. **(C)** Lung congestion scores at day 8 dpi. Bars indicate mean ± SD; p-values were determined by one-way ANOVA with Dunnett’s post hoc test comparing each treatment to PBS. **(D)** Lung viral titers measured at 8 dpi by plaque assay. Each point represents an individual animal; horizontal bars indicate mean ± SD. Dotted line denotes assay limit of detection. p-values were calculated using one-way ANOVA with multiple-comparison correction.

Further to assess disease severity, gross lung pathology was scored at necropsy. Control animals showed substantial pulmonary congestion, whereas COV2-3600- and COV2-3731 mAb-treated animals showed evidently reduced pathology, with average scores reduced to baseline levels (p < 0.0001; **Figure 5C**). Similarly, viral burden of lung homogenates revealed high titers in PBS-treated and rDENV-2D22 groups, whereas with COV2-3600, COV2-3731, or rLYCoV-1404 (positive control), we noticed either reduced or below the limit of detection (LOD) viral loads in most animals (p < 0.0001; **Figure 5D**). Collectively, our mAbs demonstrated robust prophylactic efficacy and were effective at suppressing viral replication in vivo.

### Neutralization potency of germline-reverted antibodies against SARS-CoV-2 variants

To evaluate the role of COV2-3731 somatic mutations in neutralization breadth and structural adaptation to VOCs, we produced a panel of COV2-3731 germline-reverted (GR) antibodies and assessed their neutralization activity against pseudotyped viruses such as D614G, BA.2, BA.4, and BQ.1.1 viral variants. We noted that most GR antibodies exhibited reduced neutralization potency relative to wild-type COV2-3731. Whereas several GR antibodies maintained weak to moderate neutralizing activity against the D614G virus, while activity was significantly reduced or undetectable against Omicron subvariants BA.2, BA.4, and BQ.1.1. (**Figure 6 and S8**) Further single substitutions, like G26E, possibly introduce repulsive interactions with acidic residues and disturb the CDRH1-RBD ridge stabilization. The increase in IC50s across variants can likely be attributed to this conformational destabilization (**Figure 6A**). Furthermore, an aromatic contact phenylalanine at 27 to isoleucine (F27I) disrupts an essential hydrophobic packing alongside the RBD, and this alteration decreases the buried hydrophobic area at the boundary and may shift the loop (**Figure 4E**). In contrast, threonine to Isoleucine at position 28 (T28I) likely abolishes a polar interaction, thereby waning neutralization against ancestral and Omicron variants (**Figure 6A and, S8**). Introduction of a bulky aromatic ring of tyrosine in place of a serine at 31 (S31Y) could enhance hydrophobic contacts at the apex of CDRH1 and RBD interface which elucidates the higher potency in D614G background. Subtly changes like valine to methionine at position 50 (V50M) might alter hydrophobic interactions, leading to slight changes in neutralization capacity. In distinction, proline change from serine at residue 53 (S53P) rigidified CDRH2 loop, by wiping out conformational flexibility, and removal of key hydroxyl group with the change of tyrosine to phenylalanine at 58^th^ position (Y58F) could weaken hydrogen–bond–mediated specificity (**Figure 4E, 6 and S8**). Together, rigidification of loop resulted in increase of IC_50_ across the panel, which confirms critical requirements for all changes leading to stability and flexibility in turn helping to recognize antigenically distinct spikes on VOCs (**Figure 6A**).

**Figure 6.**
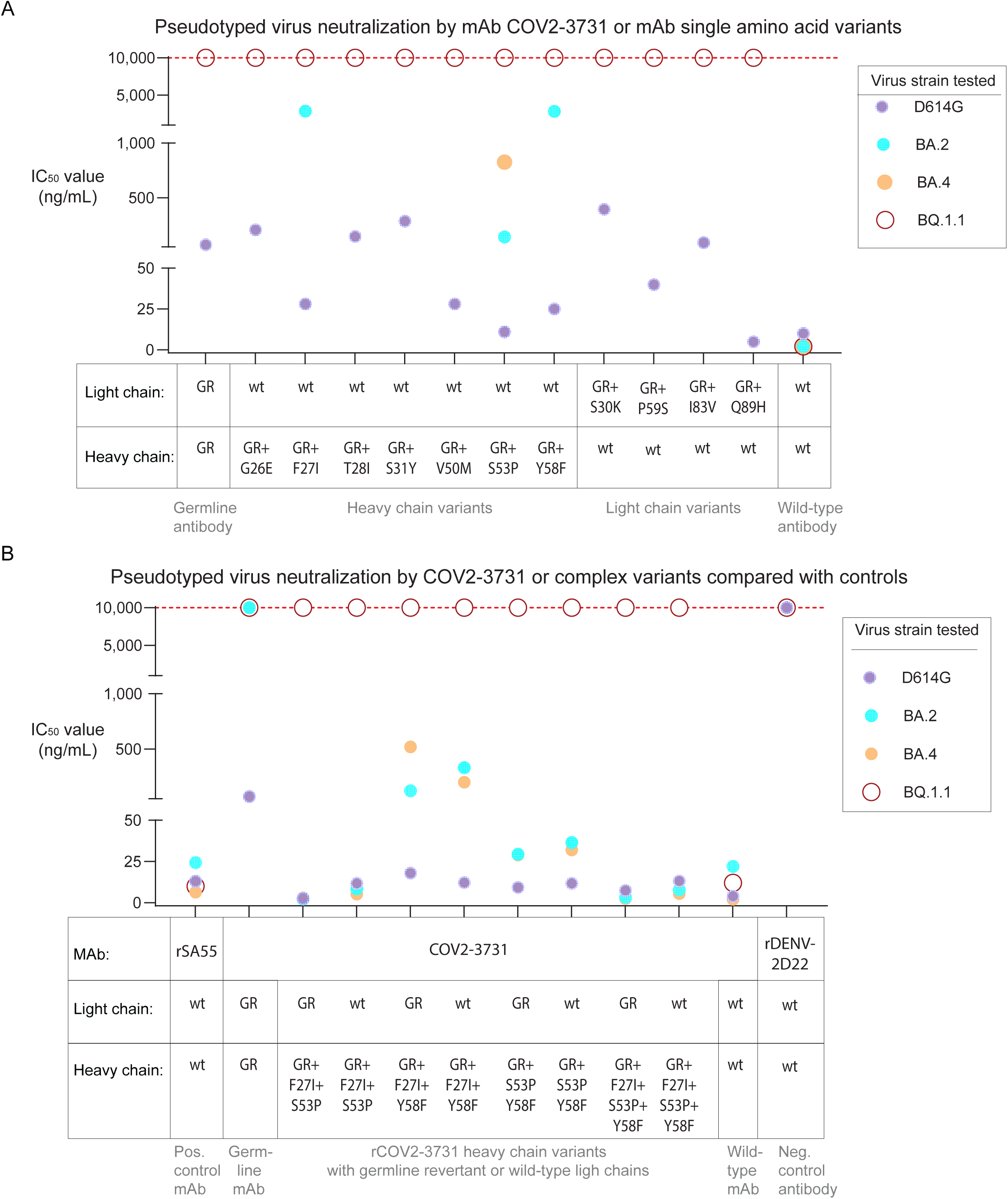
Neutralization potency of germline-reverted antibodies against SARS-CoV-2 variants. (A) Pseudotyped virus neutralization activity of germline-reverted antibodies carrying heavy or light chain substitutions. IC_50_ values (ng/mL) are shown for each monoclonal antibody (mAb) tested against pseudotyped viruses bearing Spike proteins from D614G (purple), BA.2 (cyan), BA.4 (orange), or BQ.1.1 (open red circles). Dashed red line denotes the assay upper detection limit (IC_50_ = 10,000 ng/mL). Neutralization assays were performed in technical duplicates with at least two independent experimental replicates. (B) Pseudotyped virus neutralization activity of antibodies generated from germline-reverted variants. Each combination is indicated on the x-axis. Neutralization curves were used to determine IC_50_ values against the same set of pseudotyped viruses as in (A). rSA55 and rDENV-D22 were included as positive and negative controls, respectively. Neutralization assays were performed in technical duplicates with at least two independent experimental replicates.

Analogously, light-chain substitutions generated weaker but beneficial effects, such as lysine instead of serine at the 30th position, which reconfigured CDRL1, producing variant-dependent results, while the proline to serine change at 59 (P59S) in CDRL2 likely increased flexibility that in turn partially balances for restrictive heavy-chain substitutions (**Figure 4F)**. Furthermore, we noted that changes such as arginine to valine at 18 (R18V) and glutamine to histidine (Q89H) affected neutralization modestly, which was expected to have an indirect role rather than affecting direct antigen interactions (**Figure 4F** and **6A**).

To further illustrate the effect of combined substitutions in both heavy (V_H_) and light (V_L_) chains on antigen recognition, COV2-3731 antibodies combining either germline and mature V_H_ or V_L_ were produced and tested in pseudotyped virus neutralization assays (**Figure 6B**). We noticed a combination of mutations in the heavy chain (V_H_, F27I+S53P) and matured V_L_ further reduces potency (**Figure 6B, S8 & S9**). Strikingly, antibodies comprising mature V_H_ paired with germline V_L_ maintained neutralizing activity, however, those COV2-3731 antibodies containing germline V_H_ paired with mature V_L_ presented significantly reduced potency. These data indicate that affinity maturation within the V_H_ plays a central role in expanding binding breadth and neutralization potency of the COV2-3731 mAb against divergent SARS-CoV-2 variants. Taken together, the substitutions we studied across both the V_H_ and V_L_ CDRs expose the genetic and structural basis for broad neutralization capacity of COV2-3731 and similar *IGHV3-53*/*IGKV1-33* encoded antibodies.

### Limitations of the study

Although the study investigates the biology of hundreds of antibodies including diverse somatic variants of the public clonotype encoded by *IGHV3-53/3-66* antibody genes, it should be noted the newly derived clones described here were isolated from a single convalescent individual.

## Discussion

In this study, we outline the structural, molecular, and functional basis for broad and potent mAbs that neutralize SARS-CoV-2 that are isolated from a convalescent donor infected with BA.1, particularly focused on antibodies encoded by *IGHV3-53/3-66* genes. *IGHV3-53/3-66* genes encode well-established public clonotypes that have been repetitively detected following SARS-CoV-2 infection or vaccination.^5,10,21^ Coherent with recent analyses of memory B-cell repertoires elicited by Omicron infections,^15,35–38^ We noticed extensive diversity in epitope specificities. Nevertheless, only a few antibodies preserved broad neutralization against BA.2, BA.4, XBB sublineages, and against JN.1, KP.3 and recent BA.3.2 variant that are highly immune-evasive variants^29–31^. These findings emphasize the vast antigenic remodeling of Omicron lineages, receptor-binding domain (RBD) ^15,25,26^. Regardless of this extensive evolution of RBD, our findings reveal that a select set of antibodies encoded by *IGHV3-53/3-66*, which include COV2-3600 and COV2-3731 from our panel, along with other mAbs such as ConBA-998, COVA2-04, and COVA2-39, retained neutralizing activity against extremely divergent variants. In particular, COV2-3731 structural and molecular analyses presented mechanistic insights into the flexibility of these public clonotypes, complementary to studies that describe a loss of potency among *IGHV3-53*-encoded antibodies after Omicron variant emergence ^15,39^.

Consistent with earlier reports and classifications of *IGHV3-53/3-66*-encoded antibodies^5,21^, our mAbs COV2-3600, and COV2-3731 also targeted the ACE2-binding pocket of the RBD. Further structural insights demonstrated that these antibodies realm a canonical Class 1 binding mode on RBD while engaging in an expanded network of contacts that involve N417, Y420, Y421, F456, N487, Y489, and Y505 residues, that are constrained by ACE2 binding requirements **(Figure S7)**^40^. Engagement of COV2-3731 core residues that are structurally conserved likely underlies the enhanced resistance to immune escape, in contrast to other antibodies encoded by *IGHV3-53,* which are reported to rely heavily on the F486–Y489 patch, a region commonly noted to mutate in the Omicron variant lineages ^41^. Parallelly, Class 3-like antibodies, such as COV2-3619, bind to the K444–G446 residues, thereby limiting neutralization breadth.

The remarkable COV2-3731 breadth is further confirmed by deep mutational scanning data, which revealed partial tolerance to substitutions at K417. This indeed distinguished COV2-3731 from canonical *IGHV3-53*-encoded public clonotypes, which typically lose binding upon mutation at this site^37,42–44^. Together, these outcomes imply that COV2-3731 represents an intermediate structural class, that bridges between the ACE2-competitive Class-1 and Class-3 antibodies footprints on RBD.

Germline-revertant analyses establish that COV2-3731 breadth is principally achieved by maturation of heavy-chain, particularly via substitutions in the CDRH1 and CDRH2 regions^17^. Somatic mutations in CDRH1 (G26E, F27I, T28I, S31Y) appear to jointly enhance hydrophobicity, while substitutions S53P allowed loop flexibility, whereas Y58F provided stability via an additional hydrogen-bond in CDRH2, thus allowing reorganization of the paratope to provide high-affinity binding to divergent spikes of VOCs. Our observations of COV2-3731 antibody are consistent with reports of *IGHV3-53*-encoded public clonotypes, illustrating convergent maturation pathways following multiple mRNA vaccinations ^5,21,31,45,46^. In contrast, light-chain maturation participates modestly, with substitutions in either CDRL1 or CDRL2 fine-tuning recognition of RBD. Overall, antibodies such as COV2-3731, encoded by *IGHV3-53/IGKV1-33*, maintain neutralization potency across variants, emphasizing the dominant role of heavy-chain maturation in determining breadth.

Lastly, protection studies with antibodies such as COV2-3600 and COV2-3731 confirm reductions in viral titers and prevent disease *in vivo*, consistent with previous reports of potent RBD-directed mAbs ^22,47^. Collectively, despite extensive viral evolution, we identified structurally conserved RBD motifs that not just remain a target for new generation antibodies but also provide a detailed insight for next-generation pan-variant SARS-CoV-2 vaccine design.

## Supporting information

Key Resources

## Data and materials availability

Further information and requests for resources and reagents should be directed to and will be fulfilled by J.E.C. Materials described in this paper are available for distribution for non-profit use using templated documents from the Association of University Technology Managers ‘Toolkit MTAs’, available at https://autm.net/surveys-and-tools/agreements/material-transfer-agreements/mta-toolkit.

**Supplementary information** is available for this paper.

## ACKNOWLEDGEMENTS

Cryo-EM data collections were conducted at the Center for Structural Biology Cryo-EM Facility at Vanderbilt University. Support for crystallography was provided by the Vanderbilt Center for Structural Biology. This work was supported by grant HR0011-18-2-0001 from the U.S. Defense Advanced Research Projects Agency (DARPA), grant R01 AI157155 from the U.S. National Institutes of Health (to J.E.C., M.S.D., and R.S.B.), and NIH contract HHSN75N93019C00073 (to B.J.D).

## AUTHOR CONTRIBUTIONS

N.S., and J.E.C. conceived the project; R.S.B., M.S.D., B.J.D., and J.E.C. obtained funding; N.S., S.J.Z., J.M.P., B.D., P.G., E.B., S.S., S.R., S.R.L., L.E.A., M.A., L.S.H., A.T., and S.K., performed laboratory experiments; D.C.N. and F.E.L. for providing critical reagents. L.J.M., performed bioinformatic analyses. R.S.B., M.S.D., R.H.C., J.D.B., E.D., B.J.D, and J.E.C. supervised the research; N.S., and J.E.C wrote the first draft of the paper. We thank Ivette A. Nuñez for helping with the initial experiments. All authors reviewed and approved the final manuscript.

## STAR Methods

### MATERIALS AND METHODS

#### Research participants

Studies involving human subjects or human-derived materials were approved by the Institutional Review Board of Vanderbilt University Medical Center. The subject was a healthy female, 28 years old in the U.S., with a prior history of three mRNA vaccinations encoding the ancestral SARS-CoV-2 spike protein sequence (from strain Wuhan-Hu-1). This subject was likely infected with the BA.1 or BA.1.1 variants of SARS-CoV-2, which dominated in the local circulation during December 2021. Five days post-exposure, the subject tested positive for a PCR test on nasal secretions for the presence of SARS-CoV-2, suffered upper respiratory illness symptoms and a mild clinical illness that was resolved on the sixth day, and recovered fully with only supportive therapy. Peripheral blood was obtained two months later, by phlebotomy after written informed consent. The subject underwent leukapheresis, and PBMCs were purified using a negative-selection magnetic enrichment kit (Stem CELL Technologies, Cat. no. 19654). PBMCs were cryopreserved and stored in a liquid nitrogen freezer until use.

#### Cell lines

An engineered NIH3T3 fibroblast line (mouse, male origin) constitutively expressing cell-surface human CD154 (CD40 ligand), secreted human B-cell activating factor (BAFF), and human IL-21 was kindly provided by Dr. Deepta Bhattacharya (University of Arizona, Tucson, AZ). HEK-293T/17 cells, a subclone cell line exhibiting epithelial morphology that was isolated from human embryonic kidney tissue, express the simian virus 40 (SV40) large T antigen and exhibit high transfectability. These cells were obtained from the American Type Culture Collection (ATCC, cat. CRL-11268). These HEK-293T/17 cells were cultured in DMEM supplemented with sodium pyruvate (Thermo Fisher Scientific, cat. 11995073), penicillin/streptomycin (Gibco, cat. 15140122), 25 mM HEPES (Gibco, cat. 15630-080), and 10% (vol/vol) Ultra-Low IgG FBS (Gibco, cat. 16250078) (DMEM+10%). Vero-furin cells were kindly provided by Dr. T. Pierson (NIAID, NIH) and have been described previously ^48^. FreeStyle 293F cells (Thermo Fisher Scientific, R79007) and Expi293F cells (Thermo Fisher Scientific, A1452) were maintained at 37°C in 8% CO_2_ in respective Medium (Thermo Fisher Scientific, A1435102). ExpiCHO cells (Thermo Fisher Scientific, A29127) were maintained at 37°C in 8% CO_2_ in ExpiCHO Expression Medium (Thermo Fisher Scientific, A2910002). Mycoplasma testing of Expi293F and ExpiCHO cultures was performed monthly using a PCR-based mycoplasma detection kit (ATCC, 301012K); all cell lines tested negative during these studies.

#### Viruses

The WA1/2020 strain with the D614G substitution was previously described^49^. The authentic viruses were passaged once on Vero-TMPRSS2 cells and described previously^50^. All viruses were subjected to next-generation sequencing to confirm substitutions.

#### Recombinant antigen expression and purification

Different strains of RBD subdomains of spike protein for instance, 328–531 residues of spike proteins with three previously identified stabilizing mutations (Y365F, F392W, and V395I) were incorporated into the RBD to enhance stability and yield^51^. were cloned into a mammalian expression vector downstream of a mu-phosphatase signal peptide and upstream of an AviTag and an 8× His tag. RBD plasmids encoding respective strains were transfected into Expi293F cells, and the expressed protein was purified by metal affinity chromatography on HisTrap Excel or TALON HP columns. (Cytiva). The purity and size of the expressed protein were assessed by SDS-PAGE.

BA.2 SARS-CoV-2 spike for electron microscopy studies was expressed by introducing the mutations of the BA.2 variant in the background of a previously described stabilized spike construct (VFLIP) ^52^. A T4 fibritin fold on domain, an 8x His tag, and a Twin Strep tag was added at C-terminal, while this construct also contains an inter-protomer disulfide bond, a shorter glycine-serine-rich linker between the S1 and S2 domains, and five proline substitutions relative to the native SARS-CoV-2 spike sequence. BA.2 spike protein was expressed by transfecting a plasmid encoding the COV2-S_VFLIP BA.2 construct into Expi293F cells. The transfected cell supernatants were collected after day 5, centrifuged to clarify and sterile filtered using a 0.2 µm filter after the addition of BioLock (IBA Biosciences). Ectodomain BA.2 trimeric spike protein was purified using Strep Trap XT columns (Cytiva) and eluted using 150 mM biotin in 1x protein buffer (100mM tris-150mM NaCL). The eluted BA.2 spike protein was further purified by size exclusion chromatography for use in cryo-electron microscopy (Cryo-EM).

#### B cell enrichment and flow cytometric cell sorting

PBMCs (1 × 10^8^ cells) were thawed in a 37°C water bath and mixed with cold Robo Sep TM buffer (Stem CELL Technologies, cat. no. 20104) and centrifuged briefly at 250 × *g*, for 5 minutes at room temperature. PBMCs cell pellet was resuspended in appropriate amount of cold RoboSep^TM^ buffer and placed on ice till B cell enrichment Using a negative-selection magnetic bead-based enrichment kit (EasySep ^TM^ Human B Cell Isolation Kit, STEMCELL Technologies, cat. no.17954) B cells were enriched according to the manufacturer’s protocol. Isolated B cells were washed twice with RoboSep^TM^ and then incubated with a master mix of phenotyping antibodies that includes anti-human CD19–Brilliant Violet 421TM (Bio Legend^®^, clone HIB19, cat. no. 302234), anti-human IgD–FITC (Bio Legend^®^, clone IA6–2, cat. no. 348206), and anti-human IgM–FITC (Bio Legend^®^, clone MHM-88, cat. no. 314506). each at a 1:20 dilution, for 45 minutes on ice. The stained B cells were washed with RoboSep^TM^ buffer (250 × *g*, for 3 minutes) twice to get rid excess phenotyping antibodies and incubated with SARS-CoV-2 strain BA.1 RBD protein, at a final concentration of 1µg/mL, on ice for another 45 minutes. Staining reactions were performed by adding biotinylated BA.1 RBD protein first, followed by brief washing to remove unbound RBD protein. The sample was then stained with recombinant human ACE2 (hACE2) with a FLAG tag (Sigma-Aldrich, cat. no. SAE0064), rat anti-FLAG tag-AF647 antibody (1:250 dilution, BioLegend^®^, clone L5, cat. no. 637315, lot B265929) along with 1:500 dilution of streptavidin-PE for an additional 30 minutes on ice. Cells were then washed briefly with RoboSep^TM^ buffer (250 × *g*, for 3 minutes) and resuspended in 500 µL of RoboSep^TM^ buffer for flow cytometric analysis using a SH800 cell sorter (Sony Biotechnology). BA.1 RBD, ACE2-reactive and BA.1 RBD but-ACE2 non-reactive B cells (1,345 and 462) were sorted into Medium A (Stem CELL Technologies, cat. no. 03801) containing penicillin and streptomycin. Flow cytometric data were analyzed with the SH800 software (Sony Biotechnology) and FlowJo version 10 (Tree Star). Sorted cells were then expanded on irradiated 3T3 feeder cells for 8-9 days as previously described^22^. On Day7 supernatants from antibody secreting cells (ASCs) were used for screening to confirm presence of BA.1 BD-specific antibodies by ELISA. ASCs were lifted from the specific wells flow-sorted to separate them from irradiated 3T3 feeder cells. Approximately 18,000, and 20,000 expanded ASCs were prepared for single-cell analysis on the Beacon^®^ optofluidic system (Bruker) and Chromium sequencing method (10X Genomics) to generate antibody-variable gene libraries, respectively.

#### Single-cell microfluidic assay selection of SARS-CoV-2 receptor binding domain (RBD)-reactive B cells

Flow-sorted ASCs were loaded onto an 11K chip Beacon^®^ optofluidic system (Bruker). First, cells were automatically imported onto OptoSelect 11k chips in a novel plasmablasts survival medium, which promotes cell viability and helps antibody secretion^53^. Next, using OEP technology thousands of ASCs as single-cells are guided into nL-volume chambers (NanoPens). ASCs reactivity was tested using 6-micron and 10-micron beads that are prepared by coupling recombinant biotinylated BA.1 RBD protein to streptavidin-coated polystyrene particles (Spherotech). These conjugated beads were mixed with AF568-labeled anti-human IgG (H+L) cross-adsorbed secondary antibodies (Thermo Fisher Scientific, cat. no. A-21090) at a 1:2500 dilution for the detection of secreted RBD-reactive antibodies. ASCs that were positive for SARS-CoV-2 RBD were detected through antibody binding to the conjugated beads and sequestration of fluorescent signal (AF568) from the secondary antibodies and identified by locating the NanoPens immediately adjacent to the fluorescent beads. Further, surrogate human ACE2-blocking was performed using an in-pen assay to select for antibodies that blocked human ACE2 binding to BA.1 RBD protein. To identify those such cells, ASCs were incubated with 10-micron BA.1 RBD-conjugated streptavidin-coated beads (Spherotech) in the NanoPen chambers to allow secreted antibodies to saturate the RBD. Then, a mixture containing recombinant human ACE2 (hACE2) with a FLAG tag (concentration, Sigma-Aldrich, cat. no. SAE0064), rat anti-FLAG tag-AF647 antibodies (1:50 dilution, BioLegend^®^, clone L5, cat. no. 637315, lot B265929), and anti-human IgG (H+L) cross-adsorbed-AF568 antibodies (1:2500 dilution, Thermo Fisher Scientific, cat. no. A-21090) was perfused throughout the OptoSelect 11k chip for diffusion into the NanoPen chambers. Cells secreting reactive antibodies were identified by locating pens with fluorescent (AF568) BA.1 RBD-protein-conjugated beads using the Beacon^®^ TRED filter cube. Simultaneously, hACE2 binding was detected (AF647) using a Cy5 filter cube. NanoPen chambers that contained fluorescent BA.1 RBD-protein-conjugated beads in both filter cubes were classified to contain B cells secreting BA.1 RBD-protein-reactive antibodies that did not block hACE2 binding at the concentrations tested. In contrast, NanoPen chambers that contained fluorescent BA.1 RBD-protein-conjugated beads in the TRED channel but not in the Cy5 channel were classified to contain B cells secreting BA.1 RBD-protein-reactive and hACE2 blocking antibodies. Cells of interest were exported from specific NanoPen chambers by OEP technology into individual wells of 96-well reverse transcription-PCR plates containing lysis buffer for antibody sequencing.

#### High-throughput antibody expression

For high-throughput screening, hundreds of antibodies were expressed by an approach previously described as microscale^33^. Microscale transfections were performed on Chinese hamster ovary (CHO) cell cultures (∼1 mL per antibody) using the Gibco ExpiCHO Expression System. In brief, synthesized DNA that encodes antibody (∼2 μg per transfection) was reconstituted in OptiPro serum-free medium (OptiPro SFM), and mixed with ExpiFectamine CHO Reagent, allowed to incubate 3-5 mins before adding to 800 µL of ExpiCHO cell cultures in 96-deep-well blocks using a ViaFlo 384 liquid handler (Integra Biosciences). The transfected blocks were moved to an orbital shaker at 1,000 r.p.m. with an orbital diameter of 3 mm and incubated at 37°C in 8% CO_2_. 18 hours after transfection, ExpiCHO enhancer and ExpiCHO feed reagents (Thermo Fisher Scientific) were added to the transfected CHO cells, following 4 days of incubation for a total of 5 days at 37°C in 8% CO_2_. On day 6, blocks containing transfected CHO cells and supernatants were centrifuged at 450 × *g* for 5 min, supernatants were carefully using a ViaFlo 384 liquid handler (Integra Biosciences) moved to microscale purification blocks containing 25 μL of settled protein G resin (GE Healthcare Life Sciences) per well. Post 2h incubation, protein G resin was washed with PBS and water using a 96-well plate manifold base (Qiagen) connected to the vacuum, and antibodies were eluted into 96-well PCR plates using 86 μL of 0.1 M sodium acetate buffer pH 2.7. Eluted solutions containing antibodies were neutralized by adding 5 μL of 5 M Tris-HCl pH 8.0, later neutralized antibodies were buffer-exchanged into PBS using zeba spin desalting plates (Thermo Fisher Scientific) and stored at 4°C until use.

#### MAbs and Fabs production and purification

cDNAs encoding mAbs of interest were synthesized (Twist Bioscience) and cloned into an IgG1 monocistronic expression vector (designated as pTwist-mCis_G1) or Fab expression vector (designated as pTwist-mCis_FAB) and used for production in mammalian cell culture. IgG1 monocistronic expression vector contains an enhanced 2A sequence and GSG linker that allows for the simultaneous expression of mAb heavy and light chain genes from a single construct upon transfection^54^. For mAb production, we performed transfection of ExpiCHO cell cultures using the Gibco ExpiCHO Expression System as described by the vendor. IgGs were purified from culture supernatants using HiTrap MabSelect SuRe (Cytiva) on a 24-column parallel protein chromatography system (Protein BioSolutions). Fabs were purified using the CaptureSelect column (Thermo Fisher Scientific). Purified antibodies were buffer-exchanged into PBS, concentrated using Amicon Ultra-4 50-kDa (IgG) or 30 kDa (Fab) centrifugal filter units (Millipore Sigma), and stored at 4°C until use. Purified mAbs used in animal studies were examined for endotoxin levels and found to be less than 30 EU per mg of IgG. Endotoxin testing was performed using the PTS201F cartridge (Charles River), with a sensitivity range of 10 to 0.1 EU/mL, and an Endosafe Nexgen-MCS instrument (Charles River).

#### ELISA dose-response binding assays

384-well microtiter plates were coated with recombinant SARS-CoV-2 RBD or spike proteins at a concentration of 2 µg/mL diluted in 1xDPBS at 4 °C overnight. Next day, plates were blocked with blocking buffer (2% non-fat dry milk [weight/vol] and 2% normal goat serum in DPBS-T) for 1 h after washing with DPBS containing 0.05% Tween-20 (DPBS-T). Test mAbs including positive and isotype controls were diluted in twelve 3-fold serial dilutions in blocking buffer starting 10 µg/mL concentration. After washing the plates to remove blocking buffer, diluted mAbs were added and incubated for 1 h. Plates with primary mAbs were washed once again before adding goat anti-human IgG conjugated with horseradish peroxidase (HRP) (Southern Biotech, cat. 2014-05, lot L2118-VG00B, 1:5,000 dilution in blocking buffer) and incubated for 1 h. Plates were washed for the final time to add a 3,3′,5,5′-tetramethylbenzidine (TMB) substrate (Thermo Fisher Scientific) to develop the signal. To stop the reaction 1M hydrochloric acid was added, and the absorbance was measured at 450 nm using a spectrophotometer (Biotek).

#### Competition-binding ELISA

384-well microtiter plates were coated with 1 μg/mL of purified SARS-CoV-2 BA.2 VFLIP ectodomain protein at 4°C overnight. Plates were blocked with blocking buffer (2% BSA in DPBS-T) for 1 h at room temperature. Unlabeled mAbs were diluted tenfold in blocking buffer and added to wells (20 μL per well). The plates were then incubated at room temperature for 1 h. Biotinylated preparations of recombinantly expressed reference mAbs rLY-CoV1404, rS309, or rCR3022 were then added to each respective mAb at 2.5 μg/mL in a volume of 5 μL per well (final concentration at 0.5 μg/mL) without prior washing of the unlabeled mAbs. Plates were then incubated for 1 h at room temperature. Plates were washed with DPBS-T and incubated with HRP-conjugated avidin (Sigma Aldrich, cat. A3151) for 1 h at room temperature. Bound mAbs were then detected by the addition of 25 μL of a TMB substrate, and the reaction was stopped by the addition of 25 μL of 1M HCl. Background signal was subtracted, and binding signal was normalized to the binding of each biotinylated reference mAb in the absence of competing mAbs. The following criteria were used to determine competition with the reference monoclonal antibodies (mAbs): less than 33% of the maximal binding signal of the reference mAb indicates full competition, 33 to 67% indicates partial competition, and greater than 67% indicates no competition.

#### Human ACE2 (hACE2) competition-binding ELISA

To measure ACE2 competition of ASCs supernatants to bind BA.2 spike protein was performed as previously described^22^. Briefly, a 384-well microtiter ELISA plate was coated with 2 μg/mL purified recombinant BA.2 SARS-CoV-2-S_VFLIP protein in a total volume of 25 μL at 4°C overnight. Next day, first washed plates with DPBS-T and then blocked with blocking buffer (2% non-fat dry milk and 2% normal goat serum in DPBS-T) for 1 h at room temperature. Post blocking, plates were washed with DPBS-T, and two-fold serial dilutions of ASC supernatants at a starting dilution of 1:50 were added to the wells (20 μL/well in blocking buffer) and incubated for 1 h at room temperature. Recombinant human ACE2 (hACE2) with a C-terminal FLAG tag (Sigma-Aldrich, cat. SAE0064) was added to wells at 2 μg/mL in a 5 μL volume of blocking buffer (final 0.4 μg/mL concentration of hACE2 following addition to each well) without washing. Plates were incubated for 40 min at ambient temperature. Plates were then washed with DPBS-T, and an HRP-conjugated anti-FLAG antibody (Sigma-Aldrich cat. A8592, 1:5,000 dilution in blocking buffer) was added to detect bound hACE2. After 1 h incubation at RT, plates were washed with DPBS-T, and signal was developed by the addition of 25 µL of TMB substrate followed by 25 µL of 1M HCl. ACE2 binding without a competing antibody served as a control. The signal obtained for hACE2 binding at each dilution of the supernatant tested was plotted using sigmoidal dose-response nonlinear regression analysis (Prism software version 8.0, GraphPad).

#### Generation of SARS-CoV-2 S-pseudotyped lentiviruses

All lentiviral pseudotyped viruses of SARS-CoV-2 variants (WA1/2020 D614G, BA.1.1529, BA.2.75.2, BA.2.10.4, BA.4, BR.2, BN.1, BJ.1, JD.1.1, BQ.1.1, HV.1, XBB, XBB.1.5, BA.2.86, JN.1, KP.2, KP.3, and BA.3.2) used in this study were generated using a published protocol^55^. Briefly, HEK-293T/17 cells respective lentiviral-based reporter pseudotyped viruses were generated by mixing 7.65 µg of a plasmid encoding a codon-optimized SARS-CoV-2 S gene with a 21 amino acid deletion (COV2-S_del21) under control of a CMV promoter, 22.5 µg of the pHAGE-CMV-Luc2-IRES-ZsGreen-W plasmid encoding the lentiviral genome (BEI Resources, cat. NR-52516), 4.95 µg each of the packaging plasmids HDM-Hgpm2 (BEI Resources, cat. NR-52517), HDM-tat1b (BEI Resources, cat. NR-52518), and pRC-CMV-Rev (BEI Resources, cat. NR-52519), to 1 mL of serum-free DMEM. Post mixing, 45 µL of BioT transfection reagent (Bioland Scientific, cat. B01-00) was added, and the mixture of plasmid-transfection reagent was gently pipetted up and down 3 times to mix the contents and incubated at room temperature for 5 min. Further, the transfection mixture was added dropwise to the flask while swirling gently. Approximately 16 to 18 h later, the medium was removed and fresh 2% FBS containing DMEM along with penicillin/streptomycin, sodium pyruvate, and 25 mM HEPES was supplemented. Nearly 48 h after transfection, the supernatants were collected from each flask, clarified by centrifugation, and filtered through a 0.2 µm filter. Stocks of pseudotyped virus aliquots were stored at -80°C and used in the assays later.

#### Lentiviral pseudotyped virus neutralization assays

Lentivirus-based pseudoviral neutralization assays were executed based on a previously described protocol^55^. One day prior to the assay, HEK-293T cells transduced to stably express human ACE2 (293T-hACE2 cells, BEI Resources NR-52511) were seeded onto 96-well tissue culture plates that had previously been coated with poly-D-lysine (Thermo Fisher Scientific, cat# A3890401) at a density of 1.25 × 10^4^ cells per well. On the day of the assay, mAbs were diluted using a four-fold dilution series (with each antibody in technical duplicate) in a 96-well polypropylene microtiter plate and incubated with pseudovirus for 1 h at 37°C in the presence of a final concentration of 5 µg/mL polybrene (EMD Millipore). After the 1 h incubation, pseudovirus-mAb mixtures were added to 293T-hACE2 monolayers. Plates were incubated at 37°C for 48 to 60 h, at which point cells were lysed using the Bright-Glo Luciferase Assay System (Promega) and luciferase activity was quantified using a CLARIOStar plate reader (BMG LabTech). The luminescence signal from wells in each plate where no pseudovirus or antibody was added was averaged and subtracted from each value, after which the percent infection of each well was determined relative to the average of pseudovirus-only control wells present in each plate. IC_50_ values were determined by nonlinear regression using Prism v.9.5 (GraphPad) using a four-parameter [inhibitor] vs response curve fit with top or bottom values constrained to 100 or 0, respectively. Each neutralization assay was repeated at least three times.

#### Authentic virus neutralization assays

Serial dilutions of mAbs at a starting concentration of 10 μg/mL were incubated with 10^2^ focus-forming units (FFU) of WA1/2020 D614G, B.1.351, BA.1.1, BA.1, BA.1.67.2, BA.2, BA.2.12.1, BA.4, BA.5 for 1 h at 37°C. Antibody-virus complexes were added to Vero-TMPRSS2 cell monolayers in 96-well plates and incubated at 37°C for 1 h. Subsequently, cells were overlaid with 1% (w/v) methylcellulose in MEM. Plates were harvested 30 (WA1/2020 D614G) or 72 (B.1.351, BA.1.1, BA.1, BA.1.67.2, BA.2, BA.2.12.1, BA.4, BA.5) hours later by removing overlays and fixed with 4% PFA in PBS for 20 min at room temperature. Plates were washed and sequentially incubated with a pool (SARS2-02, -08, -09, -10, -11, -13, -14, -17, -20, -26, -27, -28, -31, -38, -41, -42, -44, -49, -57, -62, -64, -65, -67, and -71 of anti-S murine antibodies^56^ (including cross-reactive mAbs to SARS-CoV) and HRP-conjugated goat anti-mouse IgG (Sigma cat. A8924) in PBS supplemented with 0.1% saponin and 0.1% bovine serum albumin. SARS-CoV-2-infected cell foci were visualized using TrueBlue peroxidase substrate (KPL) and quantified on an ImmunoSpot microanalyzer (Cellular Technologies).

#### Epitope mapping of antibodies by alanine scanning

Epitope mapping was performed as described previously^34^ using a SARS-CoV-2 S protein RBD shotgun mutagenesis mutation library (based on the Wuhan-Hu-1 strain sequence), made using a full-length expression construct for spike protein. 184 residues of the RBD (between S residues 335 and 526) were mutated individually to alanine, and alanine residues to serine. Mutations were confirmed by DNA sequencing, and clones were arrayed in a 384-well plate, one mutant per well. Binding of mAbs to each mutant clone in the alanine scanning library was determined, in duplicate, by high-throughput flow cytometry. Plasmids encoding each spike protein mutant were transfected into HEK-293T cells and allowed to express for 22 h. Cells were fixed in 4% (v/v) paraformaldehyde (Electron Microscopy Sciences) and permeabilized with 0.1% (w/v) saponin (Sigma-Aldrich) in PBS plus calcium and magnesium (PBS^++^) before incubation with mAbs diluted in PBS^++^, 10% normal goat serum (Sigma), and 0.1% saponin. MAb screening concentrations were determined using an independent immunofluorescence titration curve against cells expressing wild-type spike protein to ensure that signals were within the linear range of detection. Antibodies were detected using 3.75 μg/mL of Alexa-Fluor-488-conjugated secondary antibodies (Jackson ImmunoResearch Laboratories) in 10% normal goat serum with 0.1% saponin. Cells were washed three times with PBS^++^/0.1% saponin, followed by two washes in PBS, and mean cellular fluorescence was detected using a high-throughput Intellicyte iQue flow cytometer (Sartorius). Antibody reactivity against each mutant spike protein clone was calculated relative to wild-type spike protein reactivity by subtracting the signal from mock-transfected controls and normalizing the signal from wild-type spike-protein-transfected controls. Mutations within clones were identified as critical to the mAb epitope if they did not support reactivity of the test mAb but did support reactivity of other SARS-CoV-2 antibodies. This counter-screen strategy facilitates the exclusion of S protein mutants that are locally misfolded or have an expression defect.

#### Deep mutational scanning

BA.2 full-spike deep mutational scanning libraries were designed as described previously^43,57^. Two biological library replicates were used in all experiments. Antibody escape mapping experiments were performed by incubating each antibody for 45 min at 37°C with 10^6^ transcription units of each library replicate. Libraries were incubated with CoV2-3600, CoV2-3619, and CoV2-3731 antibodies at concentrations that neutralized >99% of library variants. After incubation, HEK-293T-ACE2 cells^25,57,58^ were inoculated with the virus-antibody mixture and, 12 to 15 h after inoculation, non-integrated viral genomes were recovered and sequenced. The escape rate for each variant in the library was calculated using a non-neutralizing standard, as described previously^39,58^. Mutation-level escape scores were measured using a biophysical model described in Yu *et al.,* and implemented in polyclonal package (v6.2)^59^. Analysis pipeline for deep mutational scanning experiments is available at https://github.com/dms-vep/SARS-CoV-2_Omicron_BA.2_spike_DMS_Crowe_mAbs.

#### Cryo-EM sample preparation and data collection

Before combining the spike protein and the Fabs, the spike was brought to room temperature and then combined at a molar ratio of 1:4 or 1:4:4 (Ag:Fab or Ag:Fab:Fab) and incubated for 1 h at 20°C. The mixture was then purified by gel-filtration using a Superose 6 Increase 10/300 GL column (Cytiva). EM grid for SARS-CoV-2 spike and Fabs COV2-3600/3619 or COV2-3731 complex were prepared fresh. A 2.2 µL volume of the purified mixture at a concentration of ∼0.46 mg/mL was applied to glow discharged (30 s at 25mA) grid (300 mesh 1.2/1.3, Quantifoil). The grids were blotted for 3.5 s before plunging into liquid ethane using Vitrobot MK4 (TFS) at 20°C and 100 % RH. Grids were screened on a Glacios (TFS) microscope. Data collection was done on Krios (TFS) operated at 300 keV equipped with a K3 and GIF (Gatan) DED detector using counting mode and a slit width of 20 eV. Movies were collected using EPU at a nominal magnification of 130,000×, pixel size of 0.647 Å/pixel, stage tilt of 0° and 30°, and defocus range of 0.8 to 1.8 µm. Grids were exposed at ∼1.12 e^-^/Å^2^/frame resulting in a total dose of ∼56 e^-^/Å^2^ (**Table S2**).

#### Cryo-EM data processing

Data processing was performed using Relion 4.0 beta 2^60^. Movies were preprocessed on-the-fly with Relion Motioncor2^61^ and CTFFind4^62^. Micrographs with low resolution, high astigmatism and high defocus were removed from the data set. The data set was first manually selected to generate 2D images and then automatically picked by the Relion template picker, and was subject to multiple rounds of 2D and 3D classification^61^. Good classes were selected and used for another round of auto-picking with Topaz training and Topaz picking^60,63^. The particles were extracted from a box of 600 pixels and binned to 160 pixels (pixel size of 2.426 Å/pixel). The particles were subjected to multiple rounds of 2D class averages, 3D initial map, and 3D classification without symmetry to obtain a clean homogeneous particle set. This set was re-extracted at a pixel size of 1.294 Å/ Å/pixel and was subjected to 3D auto-refinement. The data were further processed with CTF refine, polished^61,64^, and subjected to final 3D auto-refinement and post-processing, resulting in overall general resolution ∼3.35 / 3.23 Å (COV2-3600/3619 or COV2-3731). To better resolve the area of interaction between Cov2-RBD/Fab, a focus refinement was performed by signal subtraction with masking around the RBD/Fab (with particle expansion for COV2-3600/3619 - C3 symmetry). The subtracted particles were subjected to 3D classification without alignment, and selected particles were subjected to 3D auto-refinement and post-processing, resulting in ∼3.4 / 4.0Å map (COV2-3600/3619 or COV2-3731). Reported resolutions are based on the gold-standard Fourier shell correlation (FSC) of 0.143 criterion. Detailed statistics are provided in the **Figure. S3, S5**, and **Table S2**.

#### Model building and refinement

All the models were first docked to the map with Chimera or ChimeraX^65^. To improve the coordinates, the models were subjected to iterative refinement of manual building in Coot^66^ and Phenix^67,68^. The models were validated with Molprobity^69^ (**Table S2**). The EM map and model have been deposited into EMDB (EMD-75096, -75110, PDB 10EK for COV2-3731) and EMDB (EMD-75108, -75111, PDB 10EL for COV2-3600 and COV2-3619).

#### Focus reduction neutralization test

Serial dilutions of serum/plasma were incubated with 102 FFU of SARS-CoV-2 for 1 h at 37°C. The antibody-virus complexes were added to Vero E6 cell-culture monolayers in 96-well plates for 1 h at 37°C. Cells were then overlaid with 1% (w/v) methylcellulose in minimum essential medium (MEM) supplemented to contain 2% heat-inactivated FBS. Plates were fixed 30 h later by removing overlays and fixed with 4% paraformaldehyde (PFA) in PBS for 20 min at room temperature. The plates were incubated sequentially with 1 μg/mL of rCR3022 anti-S antibody or a murine anti-SARS-CoV-2 mAb, SARS2-16 (hybridoma supernatant diluted 1:6,000 to a final concentration of ∼20 ng/mL) and then HRP-conjugated goat anti-human IgG (Sigma-Aldrich, A6029) in PBS supplemented with 0.1% (w/v) saponin (Sigma) and 0.1% BSA. SARS-CoV-2-infected cell foci were visualized using TrueBlue peroxidase substrate (KPL) and quantified on an ImmunoSpot 5.0.37 Macro Analyzer (Cellular Technologies). Half maximal inhibitory concentration (IC_50_) values were determined by nonlinear regression analysis (with a variable slope) using Prism software.

#### MAb passive-transfer protection studies in mice

Animal studies were carried out in accordance with the Institutional Animal Care and Use Committee at UNC-Chapel Hill (protocol number 20-200). Twelve-month-old female BALB/c mice were treated with 200 μg of mAb or isotype-matched control mAb via intraperitoneal injection 12 h prior to viral challenge. Mice were anaesthetized with a mixture of ketamine/xylazine, inoculated intranasally with 10^5^ PFU of SARS-CoV-2 BA.5 MA (BA.5), and monitored daily for clinical signs of disease, weight loss, and mortality^70^. At the indicated times after infection, mice were euthanized via isoflurane overdose, and the inferior lung lobe was collected in PBS with glass beads and stored at -80°C for viral titer determination via plaque assay^71^.

#### Reverse Genetics Platform Design of SARS-CoV-2 BA.5 MA

Reverse genetics were utilized to recover a mouse adapted virus bearing the authentic mutations present in the Omicron BA.5 spike gene as previously described^70^. Briefly, using our previously described reverse genetics system for SARS-CoV-2 MA10 virus we replaced the spike gene with the variant spike gene sequence from Omicron BA.5. The virus contains a marker mutation at position T15102A to remove a naturally occurring SacI restriction enzyme site to distinguish from naturally circulating variants. Viruses were derived following systematic cDNA assembly of the infectious clone fragments, followed by *in vitro* transcription with mMessage Machine T7 polymerase (ThermoFisher, AM1344). An additional T7 reaction was set up in order to produce SARS-CoV-2 nucleocapsid mRNA. After 5 hours, the resultant mRNA was mixed with 800 μL of Vero-E6 TMPRSS2/ACE2 cells at a concentration of 10^7^ cells/mL in Opti-MEM (Gibco, 31985062). The mixture of cells and mRNA were electroporated under the following conditions: 450 Vs, 50 microfarads, 4 pulses and allowed to recover for 10 min before plating into a T75 flask with 10 mL of Dulbecco’s modified Eagle’s medium (DMEM; Gibco, 11965092) with Fetal Bovine Serum (FBS, Cytiva HyClone, 87-303HI), and 1X antibiotic/antimycotic (Gibco, 15240062). Cells were grown under 5 % CO_2_ at 37 °C and checked for CPE at 24 h post-infection (hpi). Viruses were harvested when ∼75 % of the cells in the flask demonstrated cytopathic effects (CPE), and passage 0 stocks (p0) were centrifuged at 500xg and stored at −80 °C in 1 ml aliquots. To make working stocks, 1 mL of p0 virus was inoculated into a ∼90% confluent T175 flask seeded with Vero E6 cells overexpressing human TMPRSS2/ACE2 in 24 mL of culture media. Virus supernatants were harvested when ∼75% of the cells demonstrated CPE, after which the infectious media were spun down, aliquoted, and frozen at −80 °C.

The viral titer of the stock was determined via plaque assay, whereby the virus was serially diluted ten-fold and inoculated onto confluent monolayers of Vero E6 cells in 6-well plates. Plates were incubated for 1 hour with rocking every 15 min. Subsequently, DMEM with FBS, 1X antibiotic/antimycotic, and 0.8 % agarose (Lonza, SeaKem LE Agarose, 50004) was applied as an overlay. Plaques were visualized at 3 days post-infection via staining with neutral red dye (Fisher, N129-25). Virus genomic integrity and sequence were confirmed by deep-sequencing cDNA from viral RNA on an Illumina® MiSeq sequencer.

#### Bioinformatic analysis of antibody sequences

Following sequencing, reads were demultiplexed and processed using Cell Ranger (10X Genomics, v6.1.2). In the first phase, paired heavy- and light-chain variable gene cDNA sequences containing a single productive chain pair were analyzed using PyIR^72^. Sequences were retained if they (1) lacked stop codons, (2) encoded an intact CDR3, and (3) contained an in-frame junctional region. Redundant sequences with identical amino acid sequences collapsed, and antibody variants assigned as IgM isotype by Cell Ranger V(D)J (v6.1.2) were excluded. Variable gene assignments, CDR boundaries, and somatic mutations relative to inferred germline segments were determined using PyIR.

In the second phase, paired sequence clustering was performed to define clonal families based on genetic similarity. Sequences were first grouped by identical heavy-chain V and J gene usage and HCDR3 length and then clustered at 80% nucleotide identity across the HCDR3 using a single-linkage algorithm (SciPy). Within these clusters, sequences were further partitioned by identical light-chain V and J gene usage and LCDR3 length, followed by an additional 80% identity clustering on the LCDR3. The resulting clusters were designated as clonal families. From each clonal family, the most somatically mutated sequence, as determined by PyIR, was selected for synthesis and expression. When multiple members were present, the sequence closest to the consensus was also selected.

#### Statistical analysis

For animal protection studies, a nonparametric Kruskal-Wallis test with Dunn’s post hoc correction for multiple comparisons was used to compare experimental conditions and the isotype-matched negative control. Associated *P* values are reported without assigning thresholds for statistical significance.

#### Ethics and containment procedures

All recombinant viruses were approved by the University of North Carolina at Chapel Hill Institutional Review Board under Schedule G 154002. All animal work was approved by Institutional Animal Care and Use Committee at University of North Carolina at Chapel Hill under protocol 20–200 according to guidelines outlined by the Association for the Assessment and Accreditation of Laboratory Animal Care and the U.S. Department of Agriculture. Recombinant virus proposals were reviewed by the UNC IRB and virus studies were performed in animal biosafety level 3 facilities wearing PAPR, Tyvek suits, Tyvek aprons and booties, and double gloves at University of North Carolina at Chapel Hill.

