## Supplemental Info for "Convergent IGHV3-53/3-66 antibodies elicited by Omicron BA.1 infection broadly neutralize emerging SARS-CoV-2 variants"

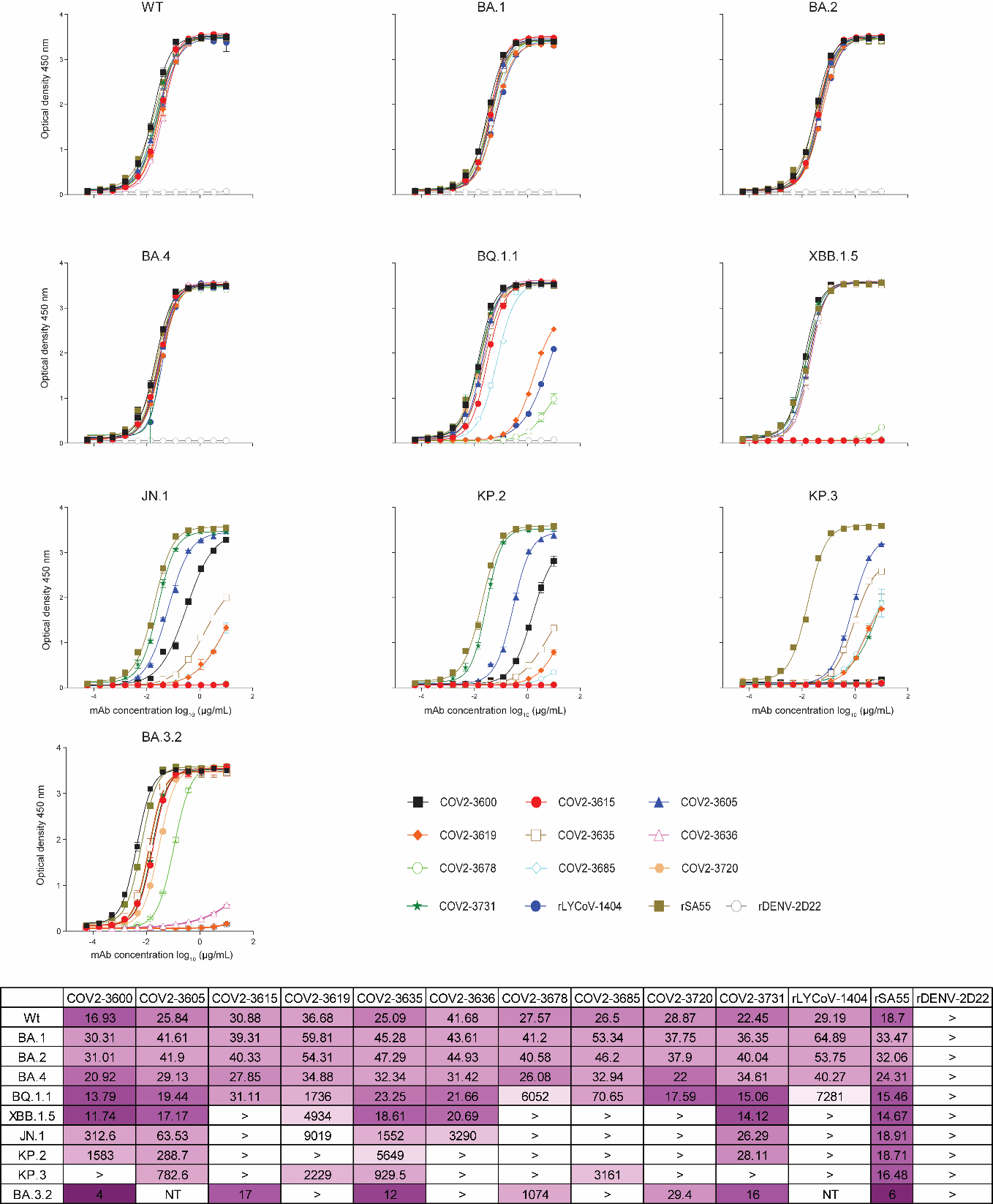

**Supplementary Figure 1**. **Binding of representative lead antibodies to diverse RBD antigens in ELISA.** ELISA binding curves are shown for a panel of monoclonal antibodies tested against the indicated antigens. Plates were coated with antigen and incubated with serially diluted antibodies. Curves show OD_450 nm_ (Y-axis) versus antibody concentration (X-axis). Each antibody is represented by a distinct color, as indicated in the key. The data shown here are representative of two independent experiments. Curves were fit using a four-parameter logistic model.

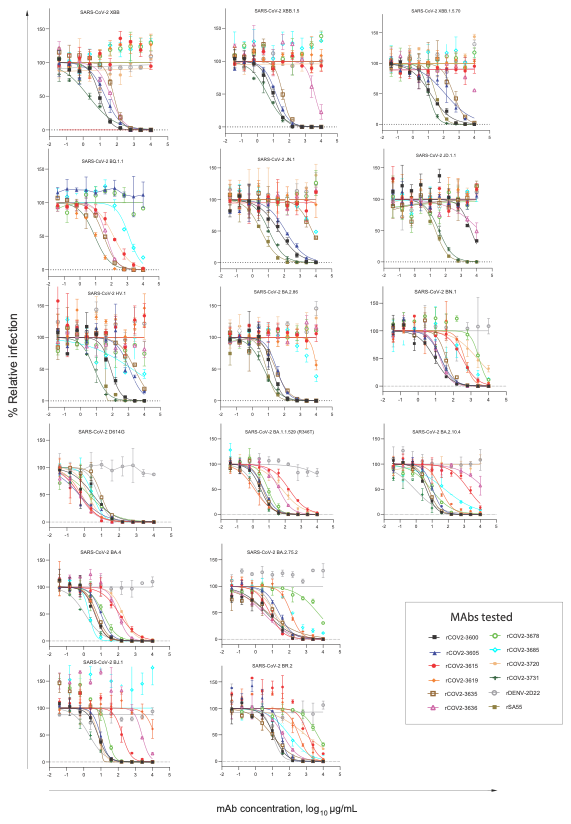

**Supplementary Figure 2**. **Neutralization breadth of lead monoclonal antibodies against SARS-CoV-2 variants measured by pseudovirus neutralization assay.** Pseudotyped lentiviral particles from ancestral SARS-CoV-2 and multiple variants of concern (VOCs) and variants of interest (VOIs), as indicated above each panel, were incubated with serial dilutions of lead monoclonal antibodies (mAbs) prior to inoculation of ACE2-expressing 293T cells. Neutralization was quantified as a percentage reduction in reporter signal relative to virus-only and cell-only controls. Each curve represents an individual monoclonal antibody (mAb) and is color-coded as shown in the key. Data points represent the mean ± error of duplicate wells from a representative experiment. Nonlinear regression curves were fit using a four-parameter logistic model.

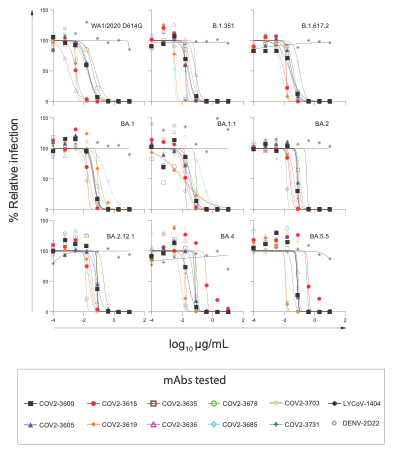

**Supplementary Figure 3**. **Neutralization breadth of lead monoclonal antibodies against SARS-CoV-2 variants measured by an authentic virus neutralization assay.** Authentic viruses were incubated with serial dilutions of lead monoclonal antibodies (mAbs) prior to inoculation of cells. Neutralization was quantified as a percentage reduction in foci relative to virus-only and cell-only controls. Each curve represents an individual monoclonal antibody (mAb) and is color-coded as shown in the key. Data points represent the mean ± error of duplicate wells from a representative experiment. Nonlinear regression curves were fit using a four-parameter logistic model.

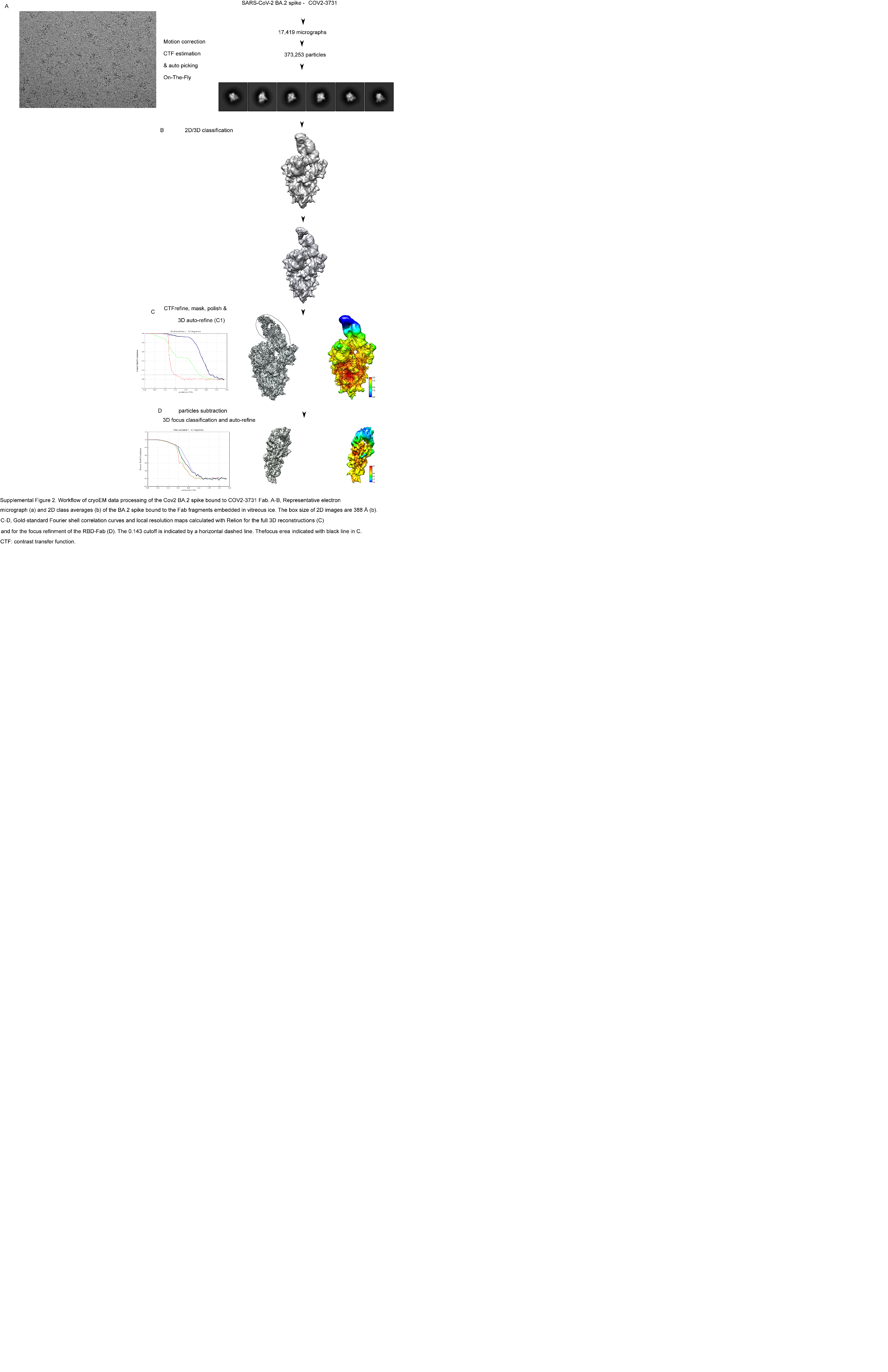

**Supplementary Figure 4. Workflow of cryoEM data processing of the SARS-CoV-2 BA.2 spike bound to COV2-3731 Fab.** **A-B**, Representative electron micrograph (**A**) and 2D class averages (**B**) of the BA.2 spike protein bound to Fab fragments embedded in vitreous ice. The box size of 2D images is 388 Å (**B**). **C-D**, Gold-standard Fourier shell correlation curves and local resolution maps calculated with Relion for the full 3D reconstructions (**C**) and for the focus refinement of the RBD-Fab (**D**). The 0.143 cutoff is indicated by a horizontal dashed line. The focus area is indicated by a black line in C. CTF: contrast transfer function.

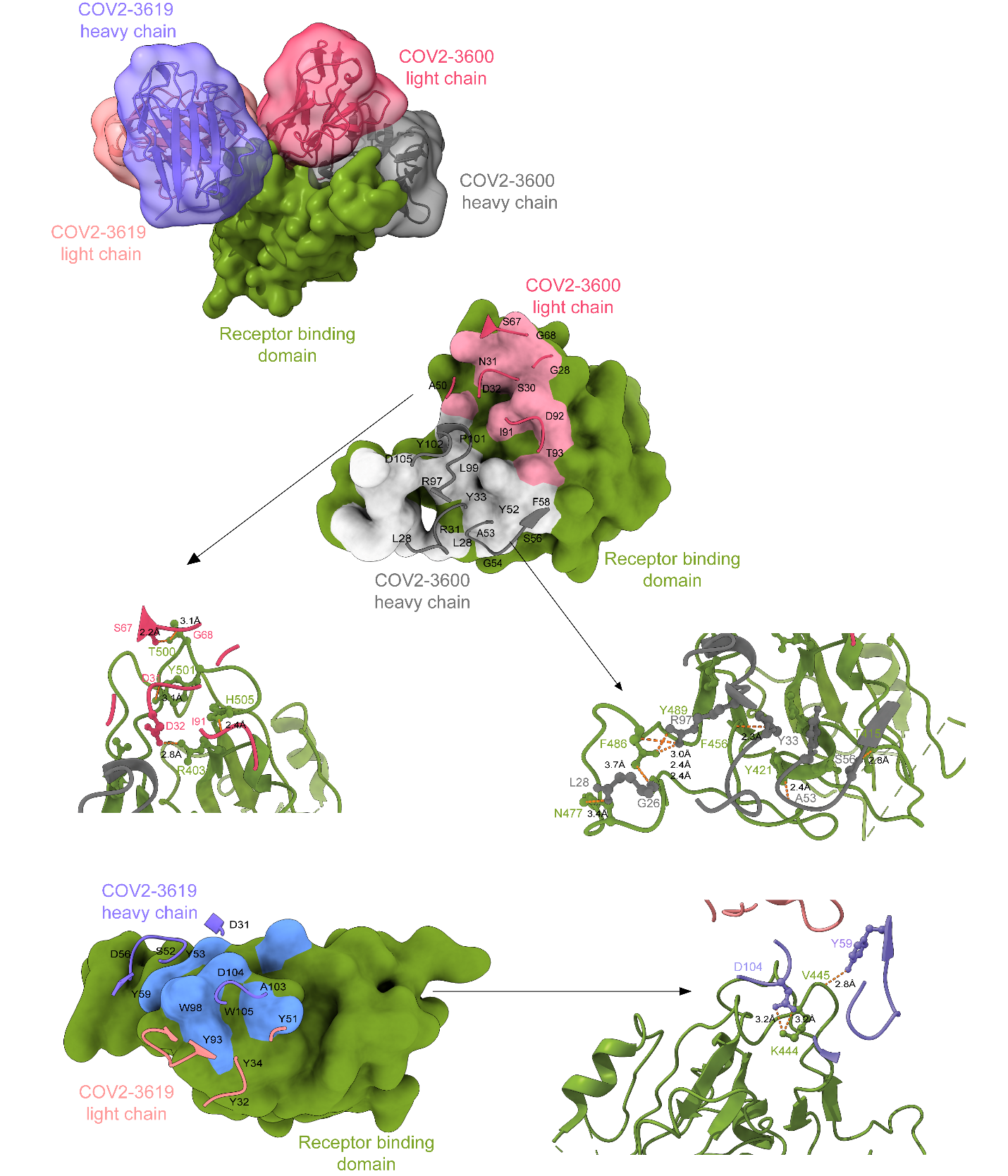

**Supplementary Figure 5. Cryo-EM structure of the COV2-3600 and -3619 Fabs in complex with BA.2 spike trimer.** (**A**) The view depicts the COV2-3600 Fab (gray and pink), COV2-3619 Fab (orange and purple) bound to one RBD (green) within the trimer (not shown). (**B**) The inset shows a zoomed-in view of the binding interface, with the heavy and light chains colored gray and pink, respectively, and the RBD in green. (**C**) Detailed view of the COV2-3600 paratope–epitope interactions, representation of RBD (green) and the COV2-3600 Fab highlighting CDR loops (heavy chain, gray; light chain CDRL1 and CDRL3, pink). (**D**) Interface details showing interactions between COV2- 3619 and the RBD.

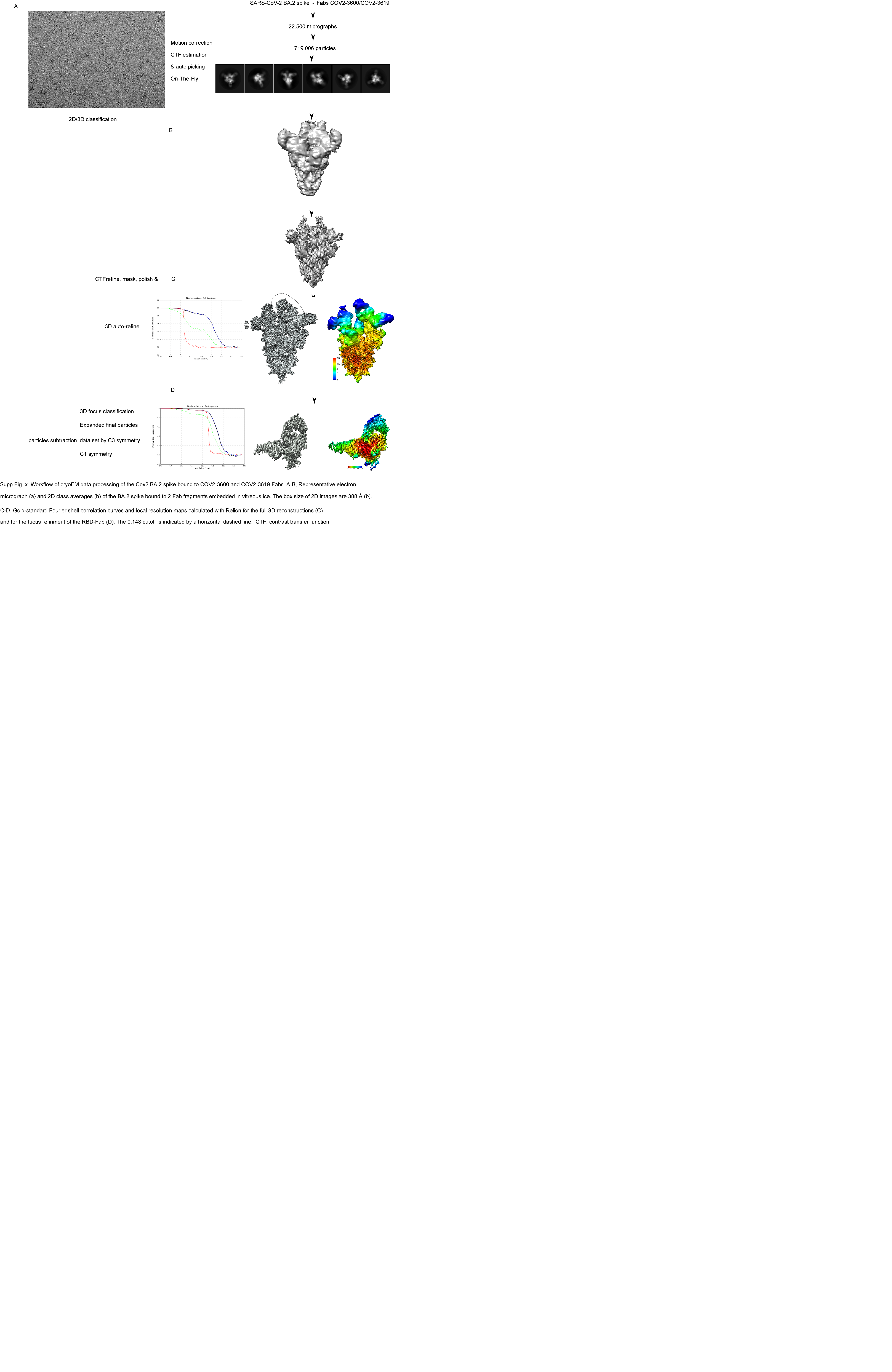

**Supplementary Figure 6**. **Workflow of cryoEM data processing of the SARS-Cov-2 BA.2 spike protein bound to COV2-3600 and COV2-3619 Fabs.** **A-B**, Representative electron micrograph (**A**) and 2D class averages (**B**) of the BA.2 spike protein bound to 2 Fab fragments embedded in vitreous ice. The box size of 2D images is 388 Å (**B**). **C-D**, Gold-standard Fourier shell correlation curves and local resolution maps calculated with Relion for the full 3D reconstructions (**C**) and for the focus refinement of the RBD-Fab (**D**). The 0.143 cutoff is indicated by a horizontal dashed line. CTF: contrast transfer function.

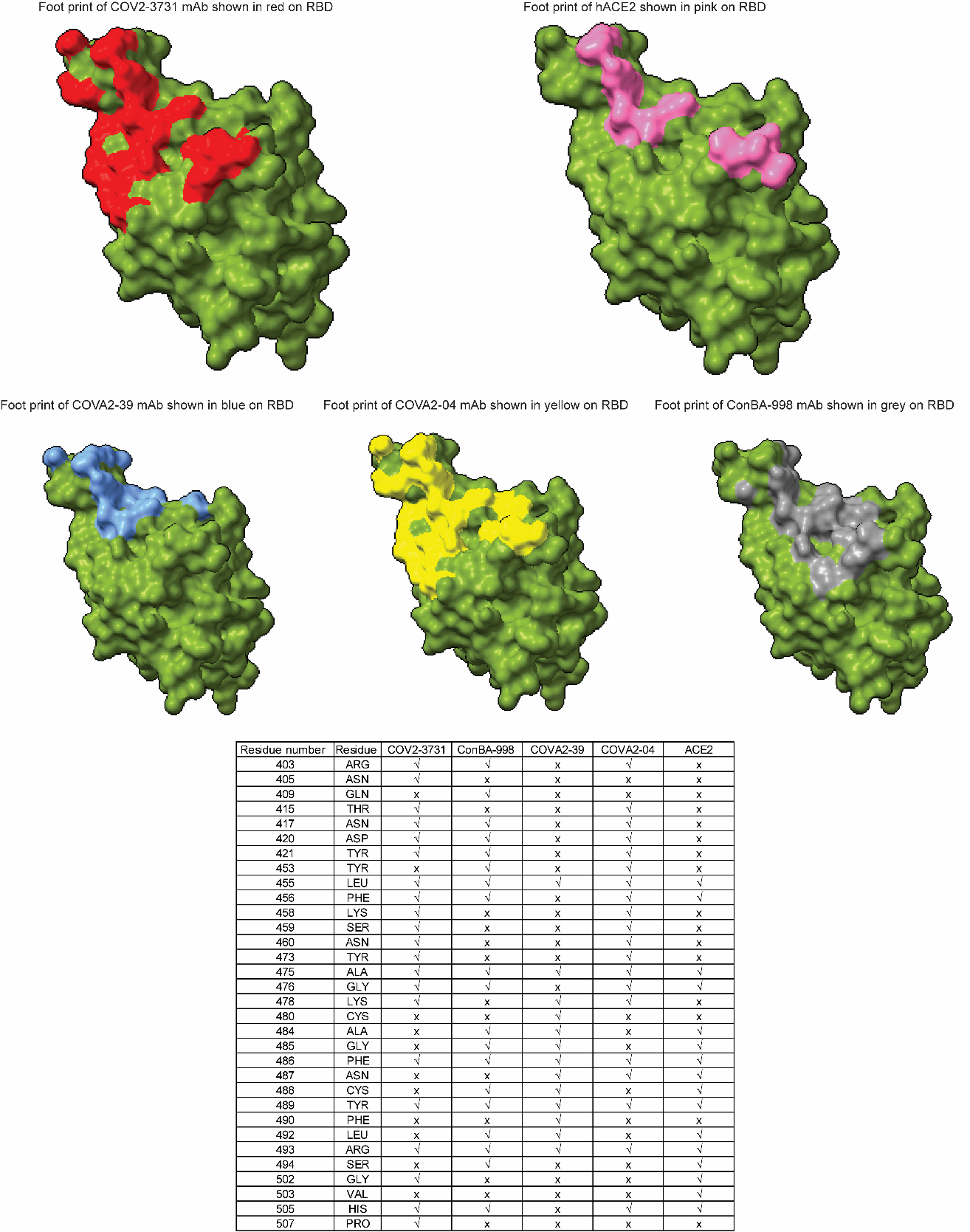

**Supplementary Figure 7.** In the top row, to the left, a detailed view of COV2-3731 Fab footprint is shown in red, and to the right, the hACE2 footprint is highlighted in pink on the RBD shown in green. In the bottom row, to the left is the detailed view of the COVA2-39 Fab footprint shown in blue; in the middle is the COVA2-04 Fab footprint shown in yellow; and to the right is the ConBA-998 Fab footprint shown in grey on the RBD shown in green. A table of residues is provided for detailed comparison of how these Fabs make contact with RBD, and see overlapping residues along with hACE2

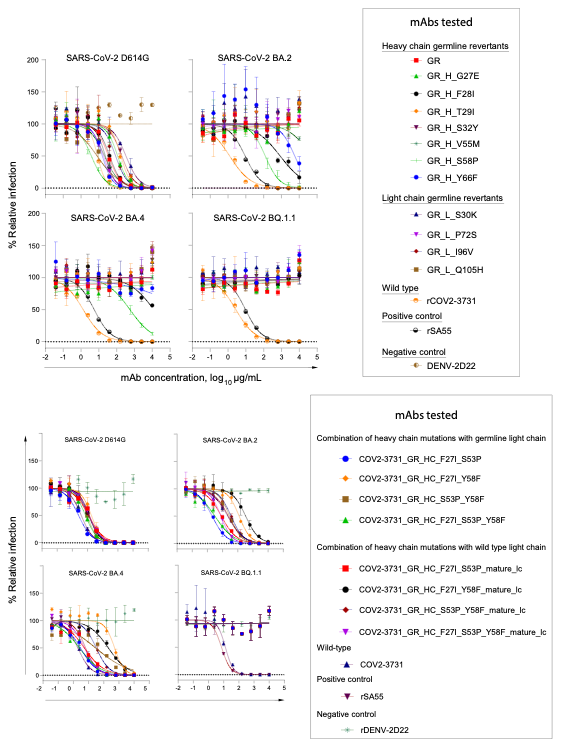

**Supplementary Figure 8**. **Neutralization breadth of COV2-3731 germline-reverted (GR) antibodies against SARS-CoV-2 variants measured by pseudovirus neutralization assay.**
Pseudotyped lentiviral particles bearing spike glycoproteins from ancestral SARS-CoV-2 and variants of concern (BA.2, BA.2 and BQ1.1) as indicated above each panel, were incubated with serial dilutions of lead monoclonal antibodies (mAbs) prior to inoculation of ACE2-expressing 293T cells. Neutralization was quantified as a percentage reduction in reporter signal relative to virus-only and cell-only controls. Each curve represents an individual monoclonal antibody (mAb) and is color-coded as shown in the key. Data points represent the mean ± error of duplicate wells from a representative experiment. Nonlinear regression curves were fit using a four-parameter logistic model.

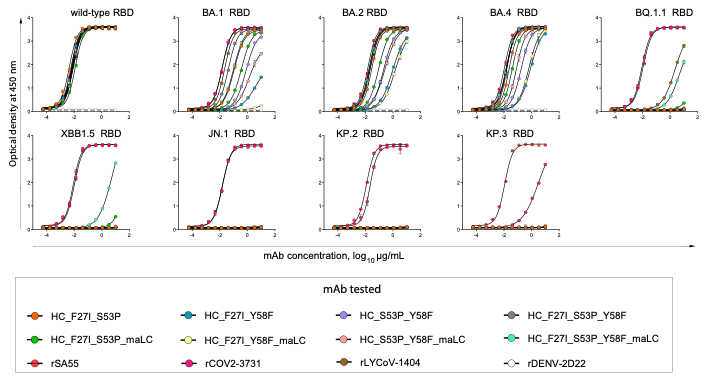

**Supplementary Figure 9. Binding of the COV2-3731 GR antibody panel to diverse RBD antigens in ELISA.** ELISA binding curves for the COV2-3731 GR antibodies as tested against the indicated antigens. Plates were coated with antigen and incubated with serially diluted antibodies. Bound IgG was detected using an HRP-conjugated secondary antibody. Curves show OD_450 nm_ (Y-axis) versus antibody concentration (X-axis). Each antibody is represented by a distinct color, as indicated in the key. The data shown here are representative of two independent experiments. Curves were fit using a four-parameter logistic model.
