## Supplementary material for "Convergent IGHV3-53/3-66 antibodies elicited by Omicron BA.1 infection broadly neutralize emerging SARS-CoV-2 variants": Key Resources

**Key resources table**

| **REAGENT or RESOURCE** | **SOURCE** | **IDENTIFIER** |
| --- | --- | --- |
| **Antibodies** | | |
| DENV r2D22 | Smith et al., 2012 | N/A |
| Goat anti-human lambda, mouse ads-UNLB | Southern Biotech | RRID:AB_2795760 |
| Goat anti-human kappa, mouse ads-UNLB | Southern Biotech | RRID:AB_2795728 |
| Goat anti-human IgG Fc, Multispecies ads-HRP | Southern Biotech | RRID:AB_2795580 |
| Goat anti-human IgG-HRP | Southern Biotech | RRID:AB_2795644 |
| **Bacterial and virus strains** | | |
| SARS-CoV-2 BA.5 MA | https://journals.asm.org/doi/10.1128/jvi.01406-25 | GenBank: [PV800150](https://www.ncbi.nlm.nih.gov/nuccore/PV800150) |
| **Biological samples** |  |  |
| Human PBMCs from leukapheresis | Vanderbilt Center for Antibody Therapeutics | N/A |
| **Chemicals, peptides, and recombinant proteins** | | |
| Avidin–peroxidase | Sigma | Cat# A3151 |
| 1-step Ultra TMB-ELISA substrate solution | Thermo Fisher | Cat# 34029 |
| ExpiCHO Expression Medium | Thermo Fisher | Cat# A2910001 |
| FreeStyle 293 expression medium | Thermo Fisher | Cat# 1238002 |
| Fetal Bovine Serum, ultra-low IgG | Thermo Fisher | Cat# 16250078 |
| EZ-Link™ NHS-PEG4-Biotin, No-Weigh™ Format | Thermo Fisher | Cat# A39259 |
| FabALACTICA^®^ Fab kit | Genovis | Cat# A2-AFK-025 |
| BioLock biotin blocking | IBA Lifesciences | Cat# 2-0205-050 |
| **Critical commercial assays** | | |
| NA |  |  |
| **Deposited data** | | |
| EMDB | EMD | EMD-75096, EMD-75110, EMD-75108 and EMD-75111 |
| PDB | PDB | 10EK (COV2-3731), 10EL (COV2-3600 and COV2-3619) |
| **Experimental models: Cell lines** | | |
| Monkey: Vero E6 +TMPRSS2 | Diamond laboratory, WUSTL | N/A |
| Monkey: Vero E6 + ACE2 + TMPRS22 | A. Creanga and B. Graham (Vaccine Research Center, NIH) | N/A |
| Monkey: Vero CCL-81 | The American Type Culture Collection (ATCC) | CCL-81; RRID: CVCL_0059 |
| Monkey: Vero Furin | Mukherjee *et al*., 2016 | N/A |
| **Experimental models: Organisms/strains** | | |
| BALB/cAnNHsd | Envigo/Inotiv | 047 |
| **Oligonucleotides** | | |
| NA |  |  |
| **Recombinant DNA** | | |
| NA |  |  |
| **Software and algorithms** | | |
| Prism | GraphPad | v 9.0.0 |
| Relion 5 | Bharat and Scheres, 2016 | [RRID:SCR_016274](https://emcore.ucsf.edu/ucsf-motioncor2) |
| CryoSPARC 4.5 | Punjani *et al*., 2017 | RRID:SCR_016501 |
| Coot 0.9.8 | Emsley *et al*., 2010 | https://www2.mrc-lmb.cam.ac.uk/personal/pemsley/coot/ |
| Phenix 1.2 | Liebschner *et al*., 2019 | https://www.phenix-online.org |
| MolProbity | Williams *et al*., 2018 | http://molprobity.biochem.duke.edu/ |
| UCSF Chimera 1.14 | Pettersen *et al*., 2004 | RRID:SCR_004097 |
| UCSF ChimeraX 1.4 |  | RRID:SCR_004097 |
| SerialEM 3.7 |  | RRID: SCR_017293 |
| Topaz 0.2.5 | Bepler *et al*., 2019, 2020 | [Topaz](https://github.com/tbepler/topaz) |
| Prism Software, version 9.1 | GraphPad Software, Inc. | https://www.graphpad.com/ |
| RTCA software, version 2.1.0 | Agilent | RTCA Software,  RRID: SCR_014821 |
| **Other** | | |
| xCELLigence RTCA MP analyzer | Acea Biosciences, Inc | N/A |
| xCELLigence E-Plate 96 PET cell culture plates | Acea Biosciences, Inc | Cat# 300601010 |
| ÄKTA pure chromatography system | GE Healthcare Life Sciences | N/A |
| AZURA chromatography system | KNAUER | N/A |
| FEI TF20 electron microscope with Gatan US4000 4k × 4k CCD camera | TFS | N/A |
| Glacios electron microscope | TFS | N/A |
| Titan Krios electron microscope | TFS | N/A |
| K3 DED | Gatan | N/A |
| Falcon3 DED | TFS | N/A |
| HisTrap Excel column | GE Healthcare Life Sciences | Cat# 17-3712-06 |
| Vitrobot MK4 | TFS | N/A |
| Superdex 200 Increase 10/300 GL Column | GE Healthcare Life Sciences | N/A |
| Quantifoil 1.2/1.3 300 mesh | Quantifoil |  |
| 400 mesh copper EM grids | EMS | Cat# 22451 |
| xCELLigence RTCA MP Analyzer | Agilent | N/A |
| xCELLigence E-Plate 96 PET cell culture plates | Agilent | Cat# 300601010 |
