## Supplementary material for "Convergent IGHV3-53/3-66 antibodies elicited by Omicron BA.1 infection broadly neutralize emerging SARS-CoV-2 variants": Suppl Data Table 2

| **Supplementary Table 2. Structural biology study features** | | | | | |
| --- | --- | --- | --- | --- | --- |
|  |  | COV2-BA.2 Fab COV-3731 | COV2-BA.2 Fab COV-3731 (focus) | COV2-BA.2 Fab COV-3600/ COV-3619 | COV2-BA.2 Fab COV-3600/ COV-3619 (focus) |
| Data Deposition | EMDB | EMD-75096 | EMD-75110 | EMD-75108 | EMD-75111 |
|  | PDB | --- | 10EK | --- | 10EL |
| Microscope setting | Microscope | Titan Krios | Titan Krios | Titan Krios | Titan Krios |
|  | Voltage (kV) | 300 | 300 | 300 | 300 |
|  | Detector | K3 | K3 | K3 | K3 |
|  | Mag | x130000 | x130000 | x130000 | x130000 |
|  | Pixel size | 0.647 | 0.647 | 0.647 | 0.647 |
|  | Exposure (e-/Å^2^) | 62.313 | 62.313 | 61.16 | 61.16 |
|  | Defocus range (μm) | 0.8-1.8 | 0.8-1.8 | 0.8-1.8 | 0.8-1.8 |
| Data | # Micrographs | 17,419 | 17,419 | 22,500 | 22,500 |
|  | # particles | 1,363,023 | --- | 1,602,611 | 2,157,018 |
|  | # particle after 2D | 1,126,366 | --- | 1,050,805 | 1,265,497 |
|  | Final particles # | 373,253 | 373,253 | 719,006 | 760,883 |
|  | Symmetry | C1 | C1 | C3 | C1 |
|  | Resolution FSC=0.143 | 3.23 | 4.1 | 3.35 | 3.4 |
| Model refinement and validation | Model resolution (Å) FSC=0.5 | --- | 4.3 | --- | 4 |
|  | Protein residues | --- | 380 | --- | 617 |
|  | Ligand | --- | --- | --- | --- |
|  | Map CC | --- | 0.62 | --- | 0.63 |
|  | RMSD |  |  |  |  |
|  | Bond lengths (Å) | --- | 0.03 | --- | 0.003 |
|  | Bond angles | --- | 0.748 | --- | 0.637 |
|  | Ramachandran |  |  |  |  |
|  | Outliers (%) | --- | 0 | --- | 0.8-1.8 |
|  | Allowed (%) | --- | 8.38 | --- | 3.98 |
|  | Favored (%) | --- | 91.62 | --- | 96.02 |
|  | Poor rotamers (%) | --- | --- | --- | 0.39 |
|  | MolProbity score | --- | 2.24 | --- | 1.93 |
|  | Clash score | --- | 19.09 | --- | 14.16 |
|  | CaBLAM score | --- | 4.83 | --- | 2.04 |
|  | B factor (Å^2^) |  |  |  |  |
|  | Protein | --- | 0.11/112.1/26.5 | --- | 0.09/259.1/34.1 |
|  | Ligand | --- | --- | --- | --- |
